# LAX28 is required for assembly of the inner dynein arm l1 and tether/tether head complex in the *Leishmania* flagellum

**DOI:** 10.1101/782888

**Authors:** Tom Beneke, Katherine Banecki, Sophia Fochler, Eva Gluenz

## Abstract

Motile eukaryotic flagella beat through coordinated activity of dynein motor proteins yet the mechanisms of dynein coordination and regulation are incompletely understood. The inner dynein arm IDA f/I1 complex and the tether/tether head (T/TH) complex are thought to be key regulators but, unlike IDA f/I1, T/TH proteins show limited conservation across flagellates. Here we characterised T/TH-associated proteins in the protist *Leishmania mexicana*. Proteome analysis of *ΔCFAP44* mutant axonemes showed that they lacked IDA f/I1 protein IC140 and a novel 28-kDA axonemal protein, LAX28. Sequence analysis identified similarities between LAX28 and the uncharacterised human sperm tail protein TEX47, sharing features with sensory BLUF-domain proteins. *Leishmania* lacking LAX28, CFAP44 or IC140 retained some motility albeit with reduced swimming speed and directionality and a propensity for flagellar curling. Expression of tagged proteins in different null mutant backgrounds showed that the axonemal localisation of LAX28 requires CFAP44 and IC140, and the axonemal localisations of CFAP44 and IC140 both depend on LAX28. These data demonstrate a role for LAX28 in motility and show mutual dependencies of IDA f/1I and T/TH-associated proteins for axonemal assembly in *Leishmania*.

**Summary Statement:** The inner dynein arm f/l1 complex is required for coordinating flagellar motility. Here we show that LAX28 is needed for its function and localization in the flagellum of *Leishmania mexicana*.

## Introduction

Eukaryotic cilia and flagella are highly conserved structures found in organism ranging from the unicellular green algae *Chlamydomonas* and protists such as *Leishmania* to specialized cells in metazoans, including humans. Even though cilia and flagella have highly conserved microtubule arrangements, structural alterations generate diversity, allowing them to exhibit different behaviours in the context of their biological function, such as signalling and motility. Dysfunctions in motile and/or non-motile cilia in humans are linked to numerous diseases collectively called ciliopathies. At least 187 ciliopathy-associated genes have been identified (Reiter and Leroux, 2017); defects in motile cilia have been shown to cause hydrocephalus in the brain, chronic respiratory problems and male infertility (Mitchison and Valente, 2017).

Motile axonemes typically have nine microtubule doublets, consisting of an A and B microtubule, and a central pair complex of two singlet microtubules. Dynein motor proteins arranged along the axoneme undergo a mechano-chemical cycle of pre- and post-power stroke conformational changes powered through the hydrolysis of ATP (Lin and Nicastro, 2018) (King, 2018). Since doublet microtubules are connected by tektin protofilaments and anchored in the basal body, conformational changes of dyneins cause the microtubules to slide against each other, resulting in flagellar bending. What remains a topic of great interest is to understand the spatio-temporal coordination of the different dynein isoforms to generate the observed flagellar waveforms (reviewed in Lindemann and Lesich (2010)).

The axoneme is organized longitudinally in regular 96 nm repeat units. Outer dynein arms (ODAs) are homogeneously spaced every 24 nm and provide the power by determining the beat frequency. Inner dynein arms (IDAs) control size and shape of the forward and reverse ciliary bend (bend amplitude). The seven subspecies of IDAs (dyneins a, b, c, d, e, f/I1 and g) are each uniquely positioned within the 96 nm repeat unit (Bui et al., 2012; Heuser et al., 2012). Additional complexes including the calmodulin and spoke associated complex (CSC) (Dymek et al., 2011), the modifier of inner arms (MIA) complex (Yamamoto et al., 2013) and the nexin-dynein regulatory complex (N-DRC) (Heuser et al., 2009; Huang et al., 1982; Oda et al., 2015; Ralston and Hill, 2006) have been shown to control the function of IDAs. Radial spokes (RSP) (Barber et al., 2012; Curry et al., 1992; Diener et al., 1993; Ralston and Hill, 2006; Williams et al., 1989; Yang et al., 2006; Yang et al., 2004) and the central pair complex (CPC) (Adams et al., 1981; Dawe et al., 2007; Dutcher et al., 1984; Lechtreck and Witman, 2007; Oda et al., 2015) influence also directly or indirectly the activity of IDAs.

Each IDA isoform is thought to have its own role in flagellar motility (Kato-Minoura et al., 1997; Kubo et al., 2018; Perrone et al., 1998). The IDA f/I1 complex has received particular attention as the centre of a regulatory hub, thought to integrate mechano-chemical signals from the CPC, RSP and other complexes (Heuser et al., 2012). IDA f/I1 is the only IDA that contains two dynein heavy chains (DHC 1α, DHC 1β) and requires an ICLC complex for incorporation into the flagellar axoneme (Heuser et al., 2012; Perrone et al., 1998; Viswanadha et al., 2014). As highlighted in Kubo et al. (2018) the *Chlamydomonas* IDA f/I1 ICLC complex contains five light chains (LC7a, LC7b, LC8, Tctex1 and Tctex2b), one accessory subunit FAP120 and two intermediate chains IC140 and IC138 (DiBella et al., 2004a; DiBella et al., 2004b; Harrison et al., 1998; Hendrickson et al., 2004; Ikeda et al., 2009; Myster et al., 1997; Myster et al., 1999; Perrone et al., 1998; Piperno, 1990; Porter et al., 1992; Smith and Sale, 1991; Toba et al., 2011). The intermediate chains have been shown to act either as regulators or assembly factors for the IDA f/I1 complex. IC140 has been shown to preassemble with both heavy chains in the cytoplasm before being transported by intraflagellar transport (IFT) proteins to the distal end of a growing flagellum (Viswanadha et al., 2014), while IC138 has been proven to be an important phosphorylation switch of IDA l1, regulating the beat amplitude, sliding velocities between microtubules and the speed of bend propagation (Hendrickson et al., 2004; VanderWaal et al., 2011).

The tether/tether head complex (T/TH) has recently emerged as a new complex linked to IDA f/I1. It was first described in *Chlamydomonas*, where cryo-electron tomography identified the T/TH structure as a link between the A-tubule and the 1*α* HC motor domain of IDA f/l1 (Heuser et al., 2012). T/TH proteins FAP44 and its paralogue FAP43 were subsequently shown to be required for assembly of the IDA f/I1 head, but not entire complex, and they are needed for regulating conformational changes of IDAs, as well as transferring their activity into microtubule sliding motion (Fu et al., 2018; Kubo et al., 2018; Urbanska et al., 2018).

FAP43 and FAP44 are part of a core group of 50 genes conserved in organisms with motile flagella (Baron et al., 2007). Their human orthologues CFAP43 and CFAP44 have been linked to non-syndromic male infertility (Krausz et al., 2015; Okutman et al., 2018; Tang et al., 2017) and loss-of-function studies confirmed a role in motility in the protists *Trypanosoma brucei* (Coutton et al., 2018) and *Leishmania mexicana* (Beneke et al., 2019). The conserved Fap57p has also been proposed to be linked to the T/TH complex (Urbanska et al., 2018). Additional components of the T/TH complex identified to date appear to be less widely conserved, such as *Chlamydomonas* FAP244 (Fu et al., 2018), MOT7, FAP102 and Cre10.g452250 (Kubo et al., 2018).

Since we found that *L. mexicana CFAP44* knockout mutants exhibited a strong motility defect, characterized by reduced swimming speed, reduced directionality (velocity/speed) and a propensity for flagellar curling (Beneke et al., 2019) we sought to identify other *Leishmania* T/TH proteins through quantitative proteomics of flagellar skeleton preparations from *CFAP44* null mutants and control cells. This identified a previously uncharacterized flagellar protein, LAX28. Further characterisation showed that LAX28, CFAP44 and the IDA f/I1 protein IC140 show mutual dependencies for localization to the flagellar axoneme and loss of LAX28 causes a similar motility defect as loss of the T/TH or IC140. Interestingly, sequence analysis identified the human protein TEX47 as a putative orthologue of LAX28, suggesting a possible role for this uncharacterised human protein in sperm motility.

## Results

### Identification of proteins missing from *ΔCFAP44* mutants axonemes

To identify new components of the *L. mexicana* T/TH complex, attempts were made to perform IP experiments with CFAP44. This was unsuccessful because CFAP44 remained associated with the axoneme following cell lysis and detergent extraction and no suitable conditions could be identified under which CFAP44 dissociated from the axoneme. CFAP44 remained associated with the axoneme in a broad range of salt concentrations, including up to 2 M LiCl, 2 M CaCl_2_, 3.2 M KCl and 4 M NaCl (Fig. S1). This stable association was then used to ask whether there were any proteins that depended on CFAP44 for their axonemal localization by comparing the protein composition of *ΔCFAP44* mutant flagella (Beneke et al., 2019) with those of the parental Cas9 T7 cell line (Beneke et al., 2017). To enrich for flagellar skeletons, NaCl-extracted flagella (Robinson and Gull, 1991) were further purified by running them over a sucrose gradient (Fig. 1 A, B and C), as this was previously shown to reduce *Leishmania* cell body contamination (Beneke et al., 2019).

**Figure 1.**
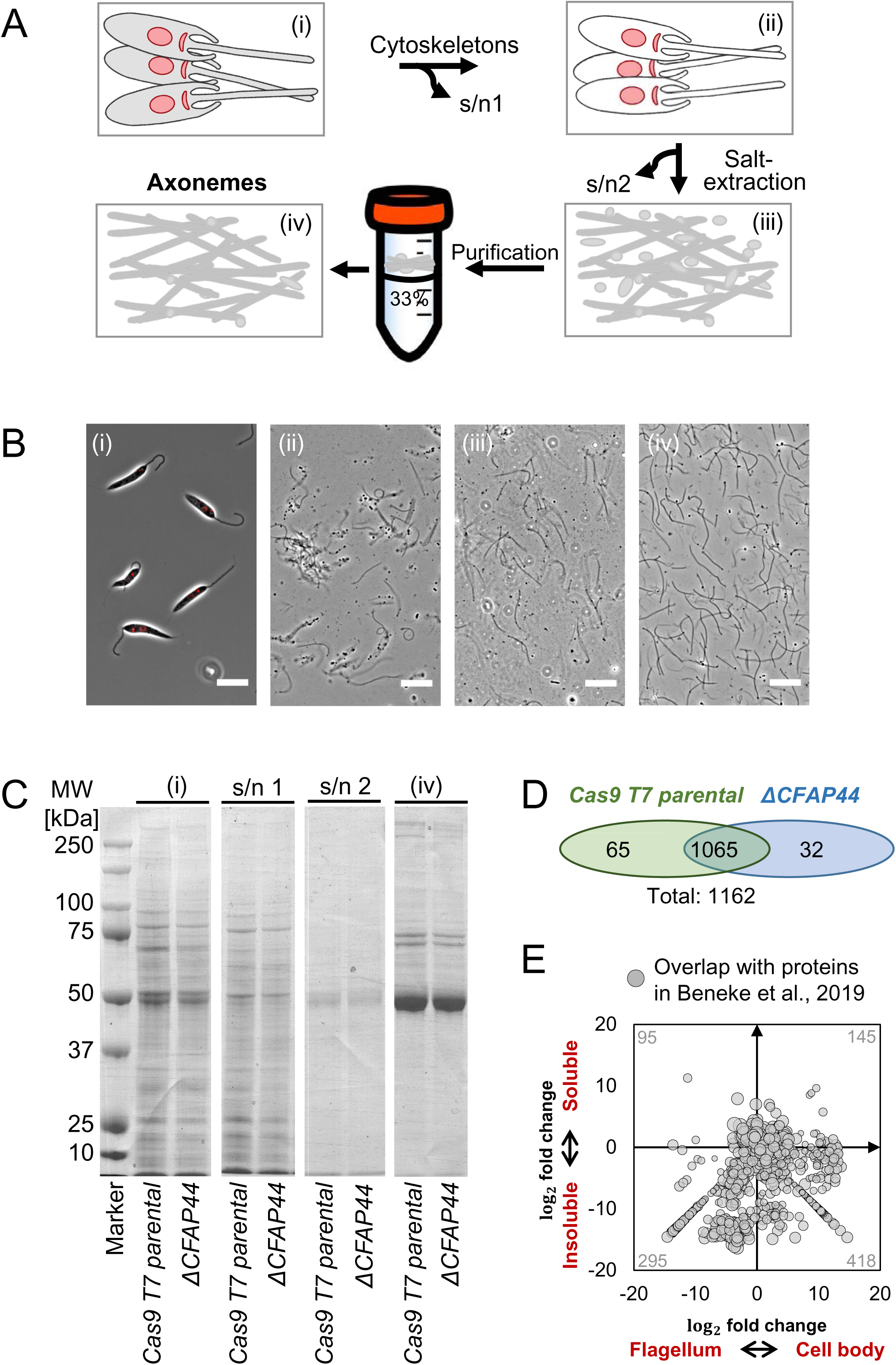
Protein composition of salt-extracted *L. mexicana* flagellar axonemes. **(A)** Overview of the flagellar isolation protocol. Percentage sucrose concentration (w/v) and isolation of supernatants (s/n) are indicated. **(B)** Micrographs show merged phase and Hoechst DNA stain (red) for each isolation stage (i-iv) depicted in (A). **(i)** *L. mexicana* cells before detergent extraction, **(ii)** cells after detergent extraction, **(iii)** axonemes after salt extraction and **(iv)** isolated salt extracted axonemes after differential centrifugation. Scale bars represent 10 μm. **(C)** Protein gel stained with Coomassie Blue. Numbers on the left indicate molecular weight in kDa. Lane **(i)** and **(s/n 1)** were loaded with protein from 4·10^6^ cells. Protein amounts for lane **(i)** and **(iv)** are as follows: *Cas9 T7 parental* **(i)** 2.02 µg and **(iv)** 0.74 µg, *ΔCFAP44* **(i)** 2.56 µg and **(iv)** 0.82 µg. 4 µg of each sample in lane **(iv)** were submitted for mass spectrometry analysis. **(D)** Venn diagram shows total number of all detected proteins (≥ 2 peptides detected, p-value > 0.95). **(E)** Proteins detected in salt extracted flagella mapped onto the SINQ enrichment plot of proteins detected in N-octyl-glucoside soluble and insoluble flagellar and cell body fractions of *L. mexicana* (Beneke et al., 2019). Each grey dot indicates a protein detected in both studies, the number of proteins is also indicated.

Liquid chromatography tandem mass spectrometry (MS) analysis of *ΔCFAP44* flagella and flagella from the *L. mex* Cas9 T7 parental cell line, for comparison (Fig. 1 C (iv)), identified a total of 1162 proteins (Fig. 1 D; S2 Table). Proteins in the mutant and parental samples overlapped well with those found in detergent-insoluble fractions of *L. mexicana*, analysed in Beneke et al. (2019) (Fig. 1 E) and included well-characterised flagellar proteins, such as CPC, N-DRC, IDA, ODA, RSP and paraflagellar rod (PFR) proteins. Whilst the mechanical method for flagellum isolation employed in Beneke et al. (2019) separated the external part of the motile flagellum form the cell body, the salt-extraction protocol preserved the connection with the basal body (Robinson and Gull, 1991). Consequently, a large number of proteins associated with the basal body, flagellar attachment zone (FAZ), intraflagellar transport (IFT) and tripartite attachment complex (TAC) were also identified (S2 Table), further expanding the inventory of *L. mexicana* flagellum-associated proteins.

To test for protein enrichment between the *ΔCFAP44* and parental flagella, a label-free normalized spectral index quantitation method SINQ (Trudgian et al., 2011) was used (S1 and S2 Table). 65 proteins were exclusively identified in the parental controls cells, while 32 proteins were exclusively identified in *ΔCFAP44* mutants.

Examination of the proteins missing from the *ΔCFAP44* flagella confirmed the loss of CFAP44 itself and its close homologue CFAP43. This is consistent with observations made for deletions and mutations on the T/TH complex in *Chlamydomonas* and *Tetrahymena* (Fu et al., 2018; Kubo et al., 2018; Urbanska et al., 2018), showing co-dependence of these proteins. Unexpectedly, the *L. mexicana* IDA f/I1 intermediate chain IC140 was also completely absent from the *ΔCFAP44* flagella. Moreover, the inner dynein arm beta and alpha heavy chains were significantly reduced in *ΔCFAP44* mutant flagella (DHC 1α -55,6%; DHC 1β -61.3%), while ODA, CP, RSP and the main component of the PFR, PFR2, showed only small changes (Table 1, S1 and S2). We confirmed depletion of IC140 from *ΔCFAP44* mutant flagella by tagging IC140 with eYFP and subsequently deleting *CFAP44* in IC140::eYFP tagged cells (Fig. 2 A). Deletion of the open reading frame (ORF) was confirmed by PCR (Fig. S2 A-F). In the tagged cell line, IC140::eYFP localized to the axoneme, as expected. By contrast, it was undetectable in *IC140::eYFP ΔCFAP44* mutants, thus confirming mass spectrometry results indicating that flagellar localisation of IC140 depends on CFAP44 in *Leishmania* (Fig. 2 A).

**Figure 2.**
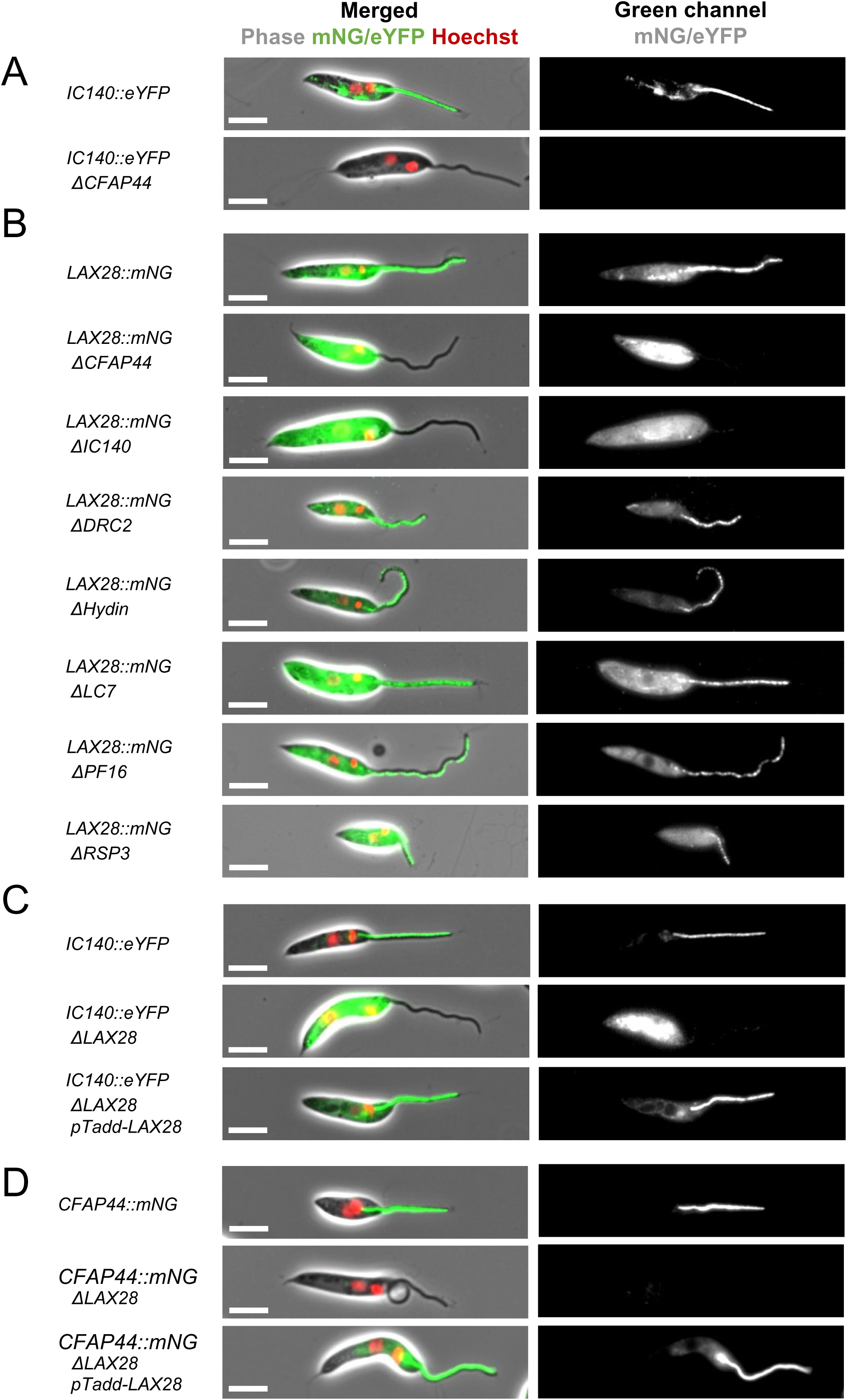
IC140 and CFAP44 are required for flagellar localization LAX28. Fluorescence micrographs showing the following *L. mexicana* cell lines: **(A)** *L. mexicana IC140::eYFP* and *L. mexicana IC140::eYFP ΔCFAP44*; **(B)** *L. mexicana LAX28::mNG* and seven different gene deletion lines, as indicated, in the *L. mexicana LAX28::mNG* background; **(C)** *L. mexicana IC140::eYFP, L. mexicana IC140::eYFP ΔLAX28::mNG*, and the addback cell line *L. mexicana IC140::eYFP ΔLAX28::mNG pTadd-LmexTEX47*; **(D)** *L. mexicana CFAP44::mNG, L. mexicana CFAP44::mNG ΔLAX28::mNG*, and the addback cell line *L. mexicana CFAP44::mNG ΔLAX28::mNG pTadd-LmexTEX47*. Left column: Merged phase and fluorescence channels; red, Hoechst-stained DNA, green, mNG or eYFP signal, respectively. Right column: grayscale rendition of green fluorescence channel. Scale bar, 5 μm.

**Table 1.**
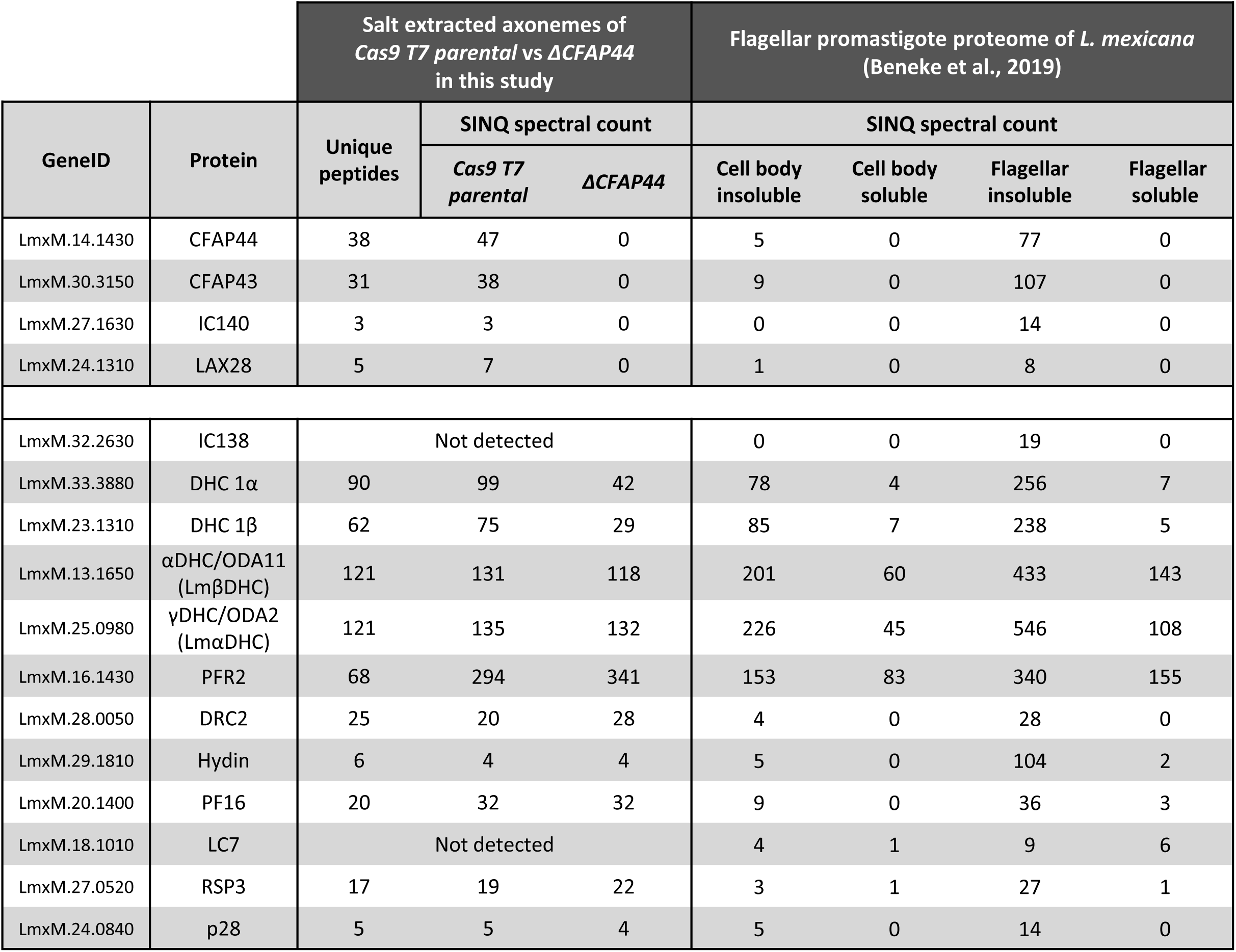
Quantitative proteomics of ΔCFAP44 mutant derived flagellar axonemes. Table shows the spectral index of SINQ quantified detected proteins in this study and in Beneke et al. (2019). Unique peptides are shown only for data from salt extracted axonemes.

We next focused on a hitherto uncharacterized protein that was completely absent from *Δ CFAP44* flagella (Table 1, S1 and S2): LmxM.24.1310 is a hypothetical protein of 254 amino acids (MW: 28,257 Da; pl: 4.9), with no defined domains, which we named LAX28 (for *<u>L</u>eishmania* <u>ax</u>onemal protein, <u>28</u> kDa). It was enriched in the *L. mexicana* flagellar proteome (Beneke et al., 2019) but had not previously been linked to T/TH or IDAs.

### Axonemal localisation of LAX28 depends on *CFAP44* and *IC140*

To provide MS-independent evidence that the flagellar localization of LAX28 is dependent on CFAP44, LAX28 was tagged with mNeonGreen (mNG) and tagged cells were then subjected to deletion of either *CFAP44*, *IC140*, *DRC2*, *Hydin*, *LC7*, *PF16* or *RSP3* (Fig. 2 B). Deletion of the open reading frame (ORF) was confirmed by PCR (Fig. S2 A, C-E). As predicted from the proteome data, LAX28::mNG localized to the axoneme of the *L. mexicana* promastigote flagellum, with some signal also in the cell body (Fig. 2B). The flagellar signal was completely lost in *ΔCFAP44* and *ΔIC140* mutants; these mutants displayed strong fluorescent signal only in the cell body and none in the flagellum. In contrast, deletion of *DRC2*, *Hydin*, *LC7*, *PF16* or *RSP3* did not alter the localization of LAX28::mNG (Fig. 2 B). This is consistent with the observed loss of the LAX28 protein from the *ΔCFAP44* flagellar protein samples (Table 1). Furthermore, it showed that loss of IC140 had the same disruptive effect on LAX28 localisation as loss of CFAP44.

### *LAX28* is required for axonemal localization of *CFAP44* and *IC140*

We next tested whether loss of LAX28 would in turn affect CFAP44 and IC140 localisation. LAX28 was deleted in cell lines expressing either *CFAP44::mNG* or *IC140::eYFP* and the deletion was confirmed by PCR as above (Fig. S2 G-I). Additionally, an addback plasmid was transfected to restore expression of LAX28 in null mutants. In the parental background, *CFAP44::mNG* and *IC140::eYFP* both localized to the axoneme (Fig. 2 C and D). Following *LAX28* deletion, the fluorescent signal was lost from the flagellum. Flagellar localization was restored in the LAX28 addback cell lines (Fig. 2 C and D). Interestingly, deletion of *LAX28* in *IC140::eYFP* cells resulted in a strong IC140::eYFP cell body signal. This was not observed for CFAP44::mNG in *ΔLAX28* mutants or IC140::eYFP in *ΔCFAP44* mutants. The biological significance of this is currently unclear. It is possible that this is a technical artefact. Alternatively, it may reflect differences in turnover of unassembled CFAP44 or IC140 proteins. Western blot analysis of the fluorescent fusion proteins (Fig. S3) showed that levels of the respective CFAP44::mNG or IC140::eYFP reporter proteins remained comparable before and after *LAX28* deletion, and following restoration of LAX28 expression (Fig. S3), indicating that the steady-state levels of CFAP44::mNG and IC140::eYFP were largely independent of LAX28.

Taken together these results suggest an essential function for LAX28 in the correct axonemal localisation of CFAP44 and IC140. Transmission electron microscopy (TEM) was used to compare the ultrastructure of *ΔLAX28* axonemes with flagella of parental cells. Multiple cross-sections of the extracellular part of the promastigote flagellum were imaged and subjected to rotational averaging (Gadelha et al., 2006; Wheeler et al., 2015). This revealed a clear lack of electron density associated with IDAs (Fig. 3 A and B), supporting the conclusion that loss of *LAX28* compromised IDA structures within the flagellar axoneme.

**Figure 3.**
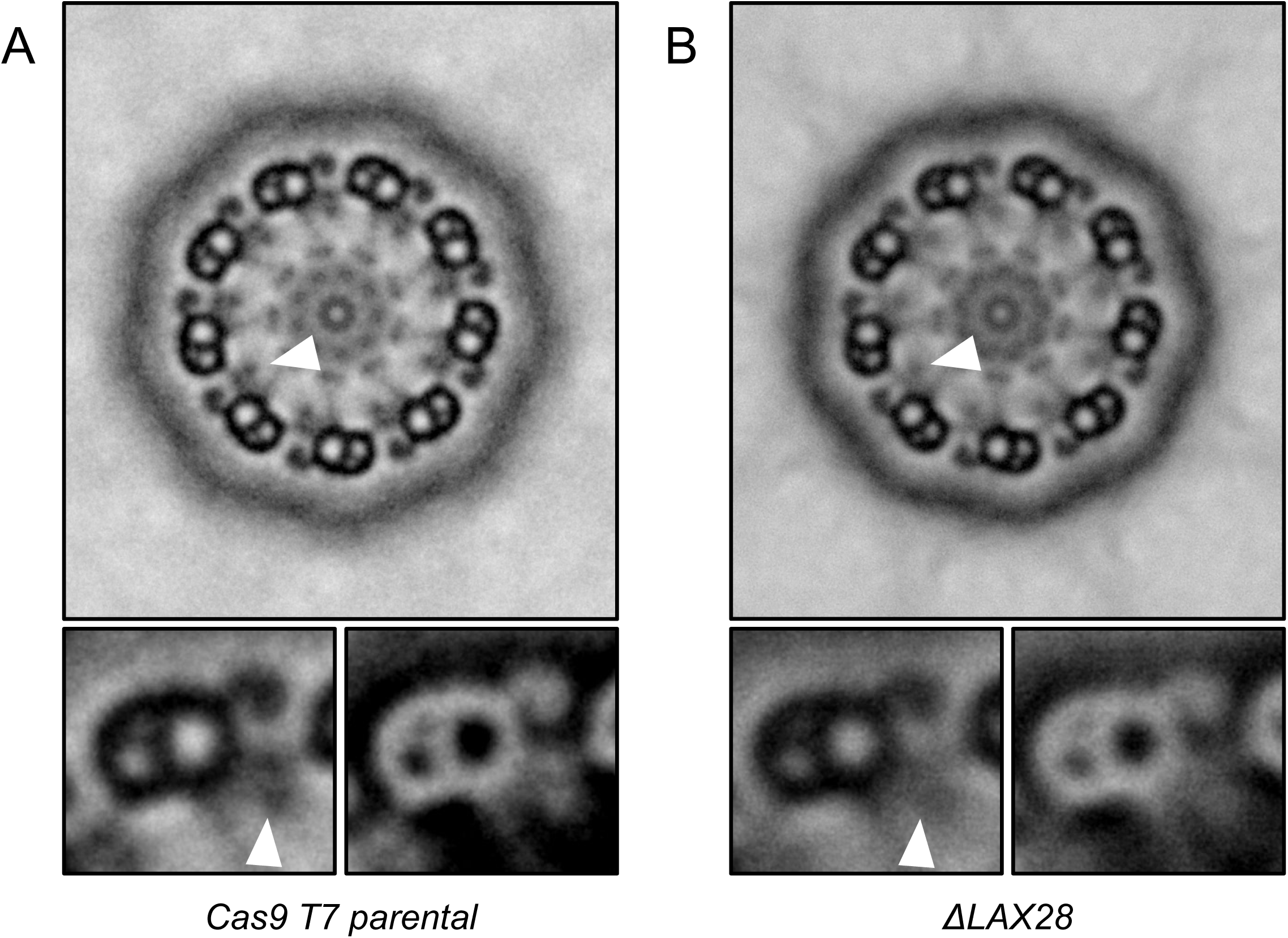
Loss of LAX28 affects IDA structure. Rotational averages of transmission electron micrographs. 9 rotational averages are stack averaged for each **(A)** *L. mex Cas9 T7 parental* cells and **(B)** *ΔLAX28* mutants. Arrows point to the position of the inner dynein arm.

This raised the question whether LAX28 is required for assembly of T/TH and IDA proteins into the axoneme, or whether the absence of LAX28 renders these structures instable following assembly. *Leishmania,* like trypanosomes, offer the opportunity to examine flagella that are being actively assembled in a cell alongside the cell’s old flagellum (Wheeler et al., 2011). The localization of LAX28::mNG, CFAP44::mNG or IC140::eYFP was imaged at different points in the cell cycle, at the end of G_1_ or the beginning of S phase (i.e. cells with 1 kinetoplast (K), 1 nucleus (N) and 1 flagellum (F)), at the beginning of new flagellum growth (1K 1N 2F) and following initiation of cytokinesis (2K 2N 2F) (Fig. 4 and S4). In the parental background, these three fusion proteins all showed a strong fluorescent signal uniformly distributed along the axonemes of old flagella and growing new flagella. Deletion of *LAX28* in CFAP44::mNG or IC140::eYFP expressing cells resulted in loss of the fusion protein from old and new flagella equally (Fig. 4). If loss of LAX28 merely rendered the T/TH and IDA structures less stable post-assembly, one might expect to find a stronger fluorescent signal in a growing new flagellum compared to an old one. The fact that this was not the case suggests that loss of LAX28 prevented stable incorporation of these proteins into the axoneme. The same results were obtained when CFAP44 or IC140 was deleted in cells expressing LAX28::mNG (Fig. S4): neither the growing nor the old flagella showed any LAX28::mNG signal. *Chlamydomonas* IC140 was previously shown to be required for assembly of the IDA f/I1 complex (Perrone et al., 1998; Viswanadha et al., 2014) and FAP44 was shown to be required for assembly of the T/TH complex (Fu et al., 2018; Kubo et al., 2018; Urbanska et al., 2018) but in these studies, loss of FAP44 did not delocalise IDA f/I1. Here we show that *Leishmania* CFAP44, IC140 and LAX28 are mutually dependent for their assembly into the axoneme.

**Figure 4.**
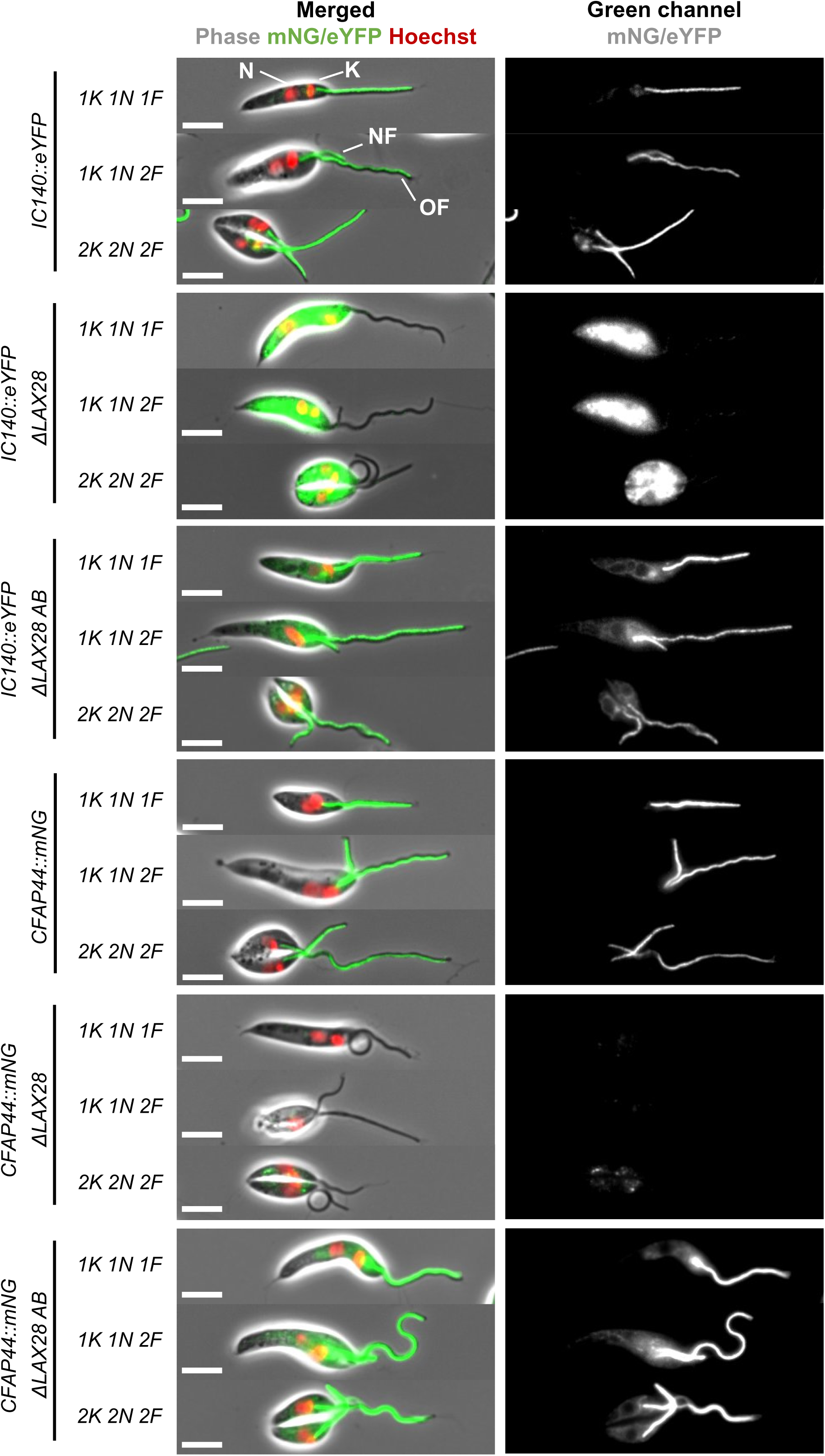
LAX28 is required for flagellar localization of IC140 and CFAP44. Fluorescence micrographs showing *L. mexicana* cell lines expressing IC140::eYFP or CFAP44::mNG reporter proteins, cell lines expressing these reporter proteins and carrying a *ΔLAX28* deletion, and deletion cell lines expressing an addbacks (AB) copy of LAX28, as indicated on the left. Three different cell cycle stages are shown for each cell line, staged according to the number of kinetoplasts (K), nuclei (N) and Flagella (F). Left column: Merged phase and fluorescence channels; red, Hoechst-stained DNA, green, mNG or eYFP signal, respectively. Right column: grayscale rendition of green fluorescence channel. Scale bar, 5 μm.

### *ΔLAX28*, *ΔCFAP44*, *ΔCFAP43* and *ΔIC140* mutants have slower swimming speeds and reduced directionality

Based on the protein localisation patterns and dependencies described above, *ΔCFAP44, Δ IC140 and ΔLAX28* mutants should exhibit similar phenotypes linked to impaired IDA f/I1 function. To test this, their respective swimming speeds and velocities were measured and the proportion of flagellar curls within the cell populations counted, comparing the characterised*Δ CFAP44*, *ΔCFAP43* and *ΔIC140* lines (Beneke et al., 2019), and newly generated*ΔLAX28* (Fig. S5) and addback cell lines. All four deletion mutants showed higher flagellar curling rates compared to the parental cell line and *ΔLAX28* addback cells (Fig. 5). The swimming speed for *ΔLAX28* was 3.8 µm/s ± 0.3 µm/s; for *ΔCFAP44* 2.9 µm/s ± 0.03 µm/s; for *ΔCFAP43* 3.3 µm/s ± 0.2 µm/s and for *ΔIC140* 2.9 µm/s ± 0.2 µm/s (Fig. 5 B and C). By contrast *ΔLAX28* addback cells showed swimming behaviours similar to parental cells (Fig. 5 B and C). In a plot of swimming speed vs. directionality, these mutants clustered closely together, showing reduced swimming speed and directionality compared to the parental controls. However, their reduction in swimming speed was not as severe as that observed in *ΔPF16* and *ΔHydin* mutants (*ΔPF16* 2.0 µm/s ± 0.1 µm/s; *ΔHydin* 2.6 µm/s ± 0.2 µm/s (Beneke et al., 2019)) and the IDA and T/TH mutants showed more directionality compared to the uncoordinated*ΔMBO2* mutants (Beneke et al., 2019) (Fig. 5B). Thus, the IDA f/I1 and T/TH mutants retained residual capacity for generation of a flagellar beat that results in displacement of cells, albeit at reduced speed.

**Figure 5.**
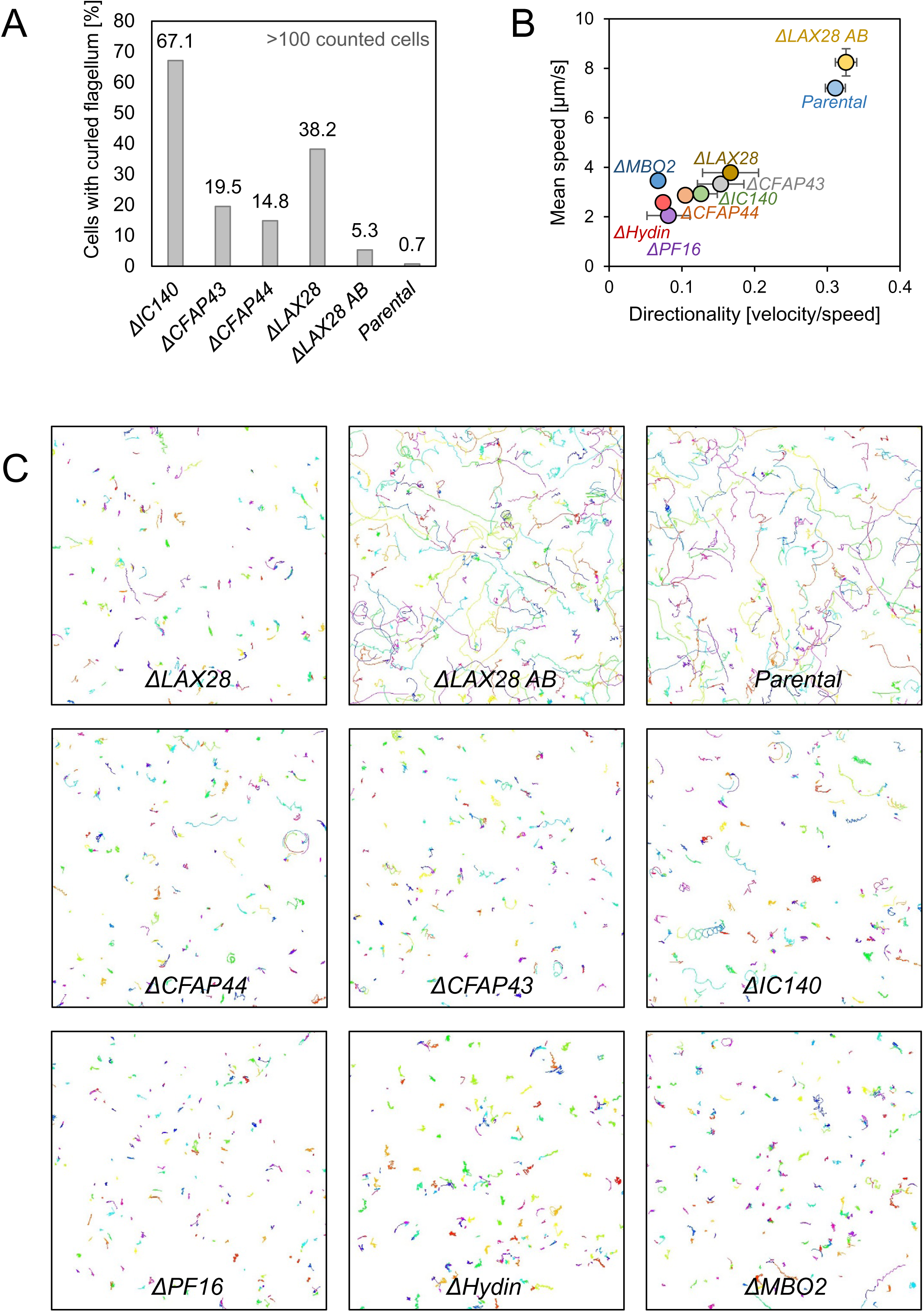
LAX28 is essential for normal flagellar motility. **(A)** Histogram showing the proportion of cells with curled flagella in *ΔIC140*, *ΔCFAP43*, *Δ CFAP44* and *ΔLAX28* mutants, the parental *L. mex Cas9 T7* cell line and *ΔLAX28* expressing a LAX28 addback copy (AB). Numbers above the bars indicate percentage of cells with curly flagella. **(B)** Plot showing mean swimming speed and directionality (the ratio of velocity to speed) for the indicated cell lines. Each point represents the average of three measurements. Error bars represent the standard deviation of the three replicates. **(C)** Swimming paths used for extracting mean swimming speed and directionality shown in (B); 200 paths are shown for each cell line.

### LAX28 shows similarity to human testis expressed protein TEX47

While LAX28 is clearly conserved across kinetoplastids, we noted a relatively low sequence identity of only 31% between the syntenic orthologs from *L. mexicana* (LmxM.24.310) and *T. brucei* (Tb927.8.6920) and initial simple protein BLAST searches did not identify any homologs outside of this lineage. A Position-Specific Iterated BLAST (PSI BLAST; (Altschul et al., 1997)) search identified the uncharacterized human Testis-expressed protein 47 (UniProt Q8TBZ9, gene TEX47 / C7orf62) as possible homologue. With a MW of 29.5 kDa, the human TEX47 protein was of a size similar to LAX28 but the sequence identity was only 16% between both proteins (Fig. 6 A). TEX47 is so named since it was found predominantly expressed in human testis (Fagerberg et al., 2014) and it was also identified in the human sperm tail proteome (Amaral et al., 2013). Interestingly, MOT7 (gene CHLRE_01g038750v5), a 26 kDa *Chlamydomonas* protein reported to interact with T/TH complex proteins (Kubo et al., 2018) also shares low similarity (24% sequence identity) with TEX47 but PSI BLAST failed to identify MOT7 when LAX28 was used as query sequence or *vice versa*. While the amino acid sequence conservation between these proteins is too low to give convincing BLAST hits, we found that LAX28, TEX47 and MOT7 are predicted by Phyre^2^ modelling (Kelley et al., 2015) to be able to adopt very similar 3D folds, as found in FAD-binding protein domains of the BLUF family (Gomelsky and Klug, 2002) (Figure 6 B).

**Figure 6.**
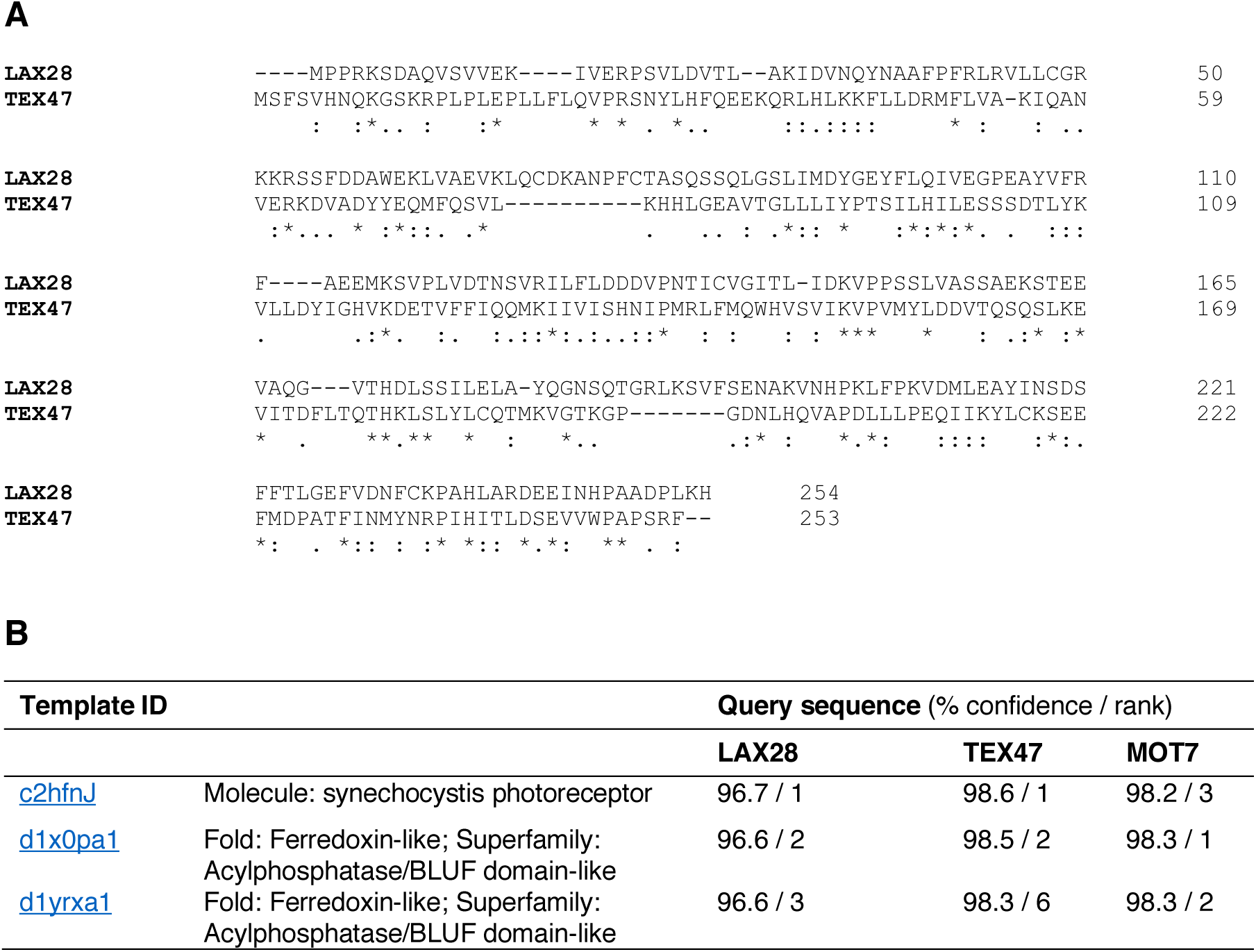
Similarity between LAX28 and human Testis-expressed protein 47. **(A)** Clustal omega protein sequence alignment (Sievers et al., 2011) of *L. mexicana* LAX28 (LmxM.24.1310) and human TEX47 (UniProt Q8TBZ9). **(B)** Results of remote homology detection and 3D fold prediction by Phyre^2^ (Kelley et al., 2015). Searches with LAX28, TEX47 or MOT7 (CHLRE_01g038750v5) all identified good matches with BLUF domain proteins, returning an overlapping set of highest-ranking hits, with high confidence scores of >96%.

**Figure 7.**
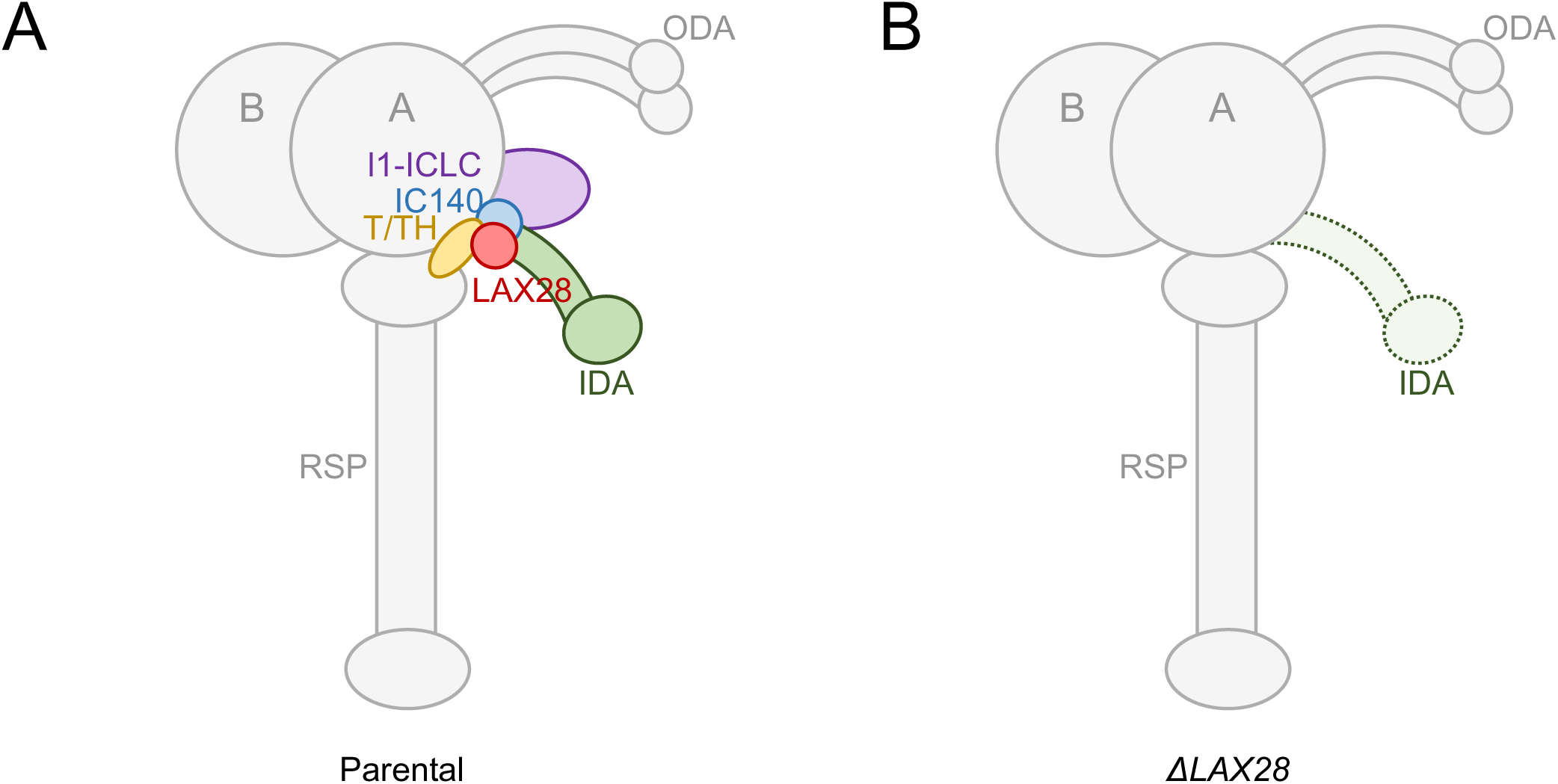
LAX28 is required for assembly of the IDA f/I1 complex within the flagellum. Cartoon showing schematic cross-sectional view of the outer microtubule doublets (A and B microtubule), radial spoke (RSP), outer dynein arms (ODA), inner dynein arms (IDA), T/TH and IDA f/I1 complex, as well as the proposed position of LAX28. **(A)** In a wild type axoneme the presence of LAX28 (red) facilitates assembly of fully functional T/TH (yellow) and IDA f/I1 complexes (violet and blue). **(B)** In *ΔLAX28* mutants the T/TH and IDA f/I1 complex are not assembled within the *Leishmania* axoneme. Both heavy chains (DHC 1α, DHC 1β) of the IDA f/I1 complex and I1-ICLC are likely to be absent. It remains unknown which, if any, of the other six IDA subspecies (dyneins a, b, c, d, e and g) are affected (green, dashed line) by the loss of LAX28.

Taken together these results show that LAX28 is essential for normal flagellar motility and indicate that the reduced flagellar motility in*ΔLAX28* knockout cells is linked to IDA f/I1 deficiency. Sequence comparisons suggest that TEX47 may have a similar function in human sperm motility.

## Discussion

### LAX28 is an inner dynein arm component required for assembly of IDA f/l1

The aim of this study was to identify T/TH complex interactors in *Leishmania*. By comparing flagellar skeleton preparations of *ΔCFAP44* null mutants and control cells, LAX28 was identified as a new potential interactor of the IDA f/I1 T/TH complex. Localisation of fluorescent fusion proteins in mutant and control cells showed clearly that LAX28 is an axonemal protein, dependent for its localization on the presence of CFAP44 and IC140. Interestingly this dependency was found to be reciprocal, as both CFAP44 and IC140 were lost from the axoneme in *ΔLAX28* mutants. One likely explanation for these observations is that LAX28 is required for incorporation of the IDA f/I1 and the T/TH complex into the 96 nm repeat. This would also explain the loss of IDA electron density observed by TEM.

Tracking *in situ* tagged IC140 and CFAP44 proteins in *ΔLAX28* mutants through the cell cycle provided supporting evidence for a role of LAX28 in IDA f/I1 assembly. In *Chlamydomonas*, IC140 binds directly to both microtubules and plays a critical role in assembly, axonemal targeting and regulation of the I1 dynein complex (Hendrickson et al., 2013). IC140 was shown to be required for assembly of both heavy chains (DHC 1α, DHC 1β) and the IDA f/I1 ICLC complex (Perrone et al., 1998; Viswanadha et al., 2014). IC140 and both heavy chains form a 20S complex and preassemble in the cytoplasm before being transported by IFT proteins to the distal end of a growing flagellum (Viswanadha et al., 2014). If a similar mechanism operates in *Leishmania*, the strong cell body signal in *IC140::eYFP* cells in the absence of *LAX28* (Fig. 2 C) may show the preassembled IDA f/I1 complex that can’t be incorporated at the tip of the growing flagellum. The observed LAX28::mNG accumulation in the cell body of *ΔIC140* or *ΔCFAP44* mutants may have a similar explanation (Fig. 2 B). This is in contrast to the complete loss of CFAP44::mNG signal following loss of *LAX28* (Fig. 2 D) and complete depletion of IC140::eYFP signal in *ΔCFAP44* mutants (Fig. 2 A). Setting aside the possibility of technical variations, this could indicate that CFAP44 is not assembled into the IDA f/I1 20S complex in the cytoplasm is or subject to more rapid turnover in cells that lack LAX28 and that IC140 is degraded more quickly in absence of CFAP44.

Loss of LAX28 reduces swimming speed and directionality to levels similar to those observed for *ΔIC140* or *ΔCFAP44* mutants, consistent with a functional link between these proteins in *Leishmania.* With respect to flagellar curling rates, the *ΔLAX28* cells (38% curling) do however not fully phenocopy the *ΔIC140* cells (67% curling). There are at least two alternative explanations that could account for this discrepancy: first, the loss of LAX28 reduced IC140 protein to levels below the detection limit of the fluorescence microscope, but it is possible that a sufficiently small number of IC140 proteins remained in the axoneme, so that curling rates in*ΔLAX28* mutants remained below those observed in an *IC140* null mutant. Alternatively, the loss of LAX28 may affect other IDA-related structures in addition to IDA f/I1, either some of the single-headed IDAs or accessory complexes. This loss may counteract the effect of IC140 loss to some extent, so that the net result is an intermediate rate of curling.

While the reduction of TEM electron density at the location of IDAs in *ΔLAX28* mutants (Fig. 3) is consistent with an IDA f/I1 assembly defect, the residual electron density indicates that at least some of the other IDA heavy chains (a-e and g) remain. This aligns with previous findings in *Chlamydomonas*, where each IDA is independently targeted to its location within the 96 nm repeat, causing gaps if assembly of one IDA fails. This gap will be reflected as reduced electron density in thin sections, but not complete absence of signal (Bui et al., 2012; Heuser et al., 2012; Piperno, 1990). In *Leishmania*, the flagella of IDA f/I1 deletion mutants are not completely paralysed but their swimming speed and directionality is reduced compared to wild type cells (Beneke et al., 2019) (Fig. 5 B and C). Similar observations have been made for *Δ IC140* mutants and IDA f/I1 T/TH complex mutants in *Chlamydomonas. ΔFAP44* and *ΔIC140* mutants showed only moderate flagellar beat and swimming defects compared to other ODA- or IDA-heavy chain deficient mutants which are more severe (Kato-Minoura et al., 1997; Kubo et al., 2018; Perrone et al., 1998). While it is clear that LAX28 is required for normal motility and function of the *Leishmania* flagellum, defining the precise IDA assembly defects and fate of the other IDA heavy chains (a-e and g) and other ultrastructural changes will require cryo-EM reconstructions of parental and *ΔLAX28* mutant axonemes.

### The IDA f/I1 and T/TH complex shows diversity across flagellated eukaryotes

Biochemical studies and cryo-EM reconstructions of the *Chlamydomonas* T/TH complex showed FAP44 to be a constituent of the tether, and demonstrated direct interactions between FAP44, FAP43 and IDA f/I1 dynein motor domains I1*α* and I1*β*. (Fu et al., 2018; Kubo et al., 2018). The tether is anchored to the axoneme through the C-terminal coiled-coil domains of FAP44 and FAP43. Ultrastructural comparisons between wild type and a *fap44* mutants suggested that the tether constrains the nucleotide-dependent movement of the IDA f/I1 head domains (Kubo et al., 2018) and indicated a role for FAP44 in the assembly of the IDA f/I1 dynein motor but not the ICLC (Fu et al., 2018). This is in contrast to the results from *L. mexicana*, where mass spectrometry data of salt extracted flagellar preparations and *in situ* tagging showed a loss of IC140 in *ΔCFAP44* mutants as well as a reduction of both IDA l1 α and β heavy chains (Table 1, Fig. 2 A). *L. mexicana* CFAP43 was also lost from *ΔCFAP44* mutant axonemes, in line with the finding that the *Chlamydomonas* T/TH complex requires dimerization of C/FAP43 and C/FAP44 to be assembled (Fu et al., 2018; Urbanska et al., 2018). Furthermore, our data show that *L. mexicana* CFAP44 and IC140 are both dependent on LAX28. These data suggest the *Leishmania* T/TH complex cannot assemble in the absence of the IDA f/I1 complex, contrary to the case in *Chlamydomonas* (Heuser et al., 2012) (Fu et al., 2018). Further more detailed studies of the *L. mexicana* wild type axoneme ultrastructure could help to define the precise location of LAX28 relative to the T/TH and IDA f/I1 complex and define the fate of the T/TH structure in a range of defined IDA f/I1 mutants.

Of the known T/TH associated proteins, CFAP44, CFAP43 and Fap57p (Urbanska et al., 2018) seem to be well conserved across flagellated eukaryotes, including *Trypanosoma*. Fap57p has been identified to be a homologue of CMF6 and CMF7 in *T. brucei* (Baron et al., 2007). A partial *CMF6* deletion mutant showed swimming patterns similar to *ΔLAX28*, *ΔCFAP44* and *ΔIC140* knockouts (Beneke et al., 2019). Other T/TH associated proteins reported to date appear to be rather restricted to the taxa in which they were found: *Chlamydomonas* proteins Cre10.g452250 (Kubo et al., 2018) and FAP244 were identified to be part of the T/TH complex but have no homologue in *Tetrahymena* and other organisms (Fu et al., 2018). Interestingly, CFAP44, CFAP43, Fap57p and FAP244 all share similar domain architectures, with N-terminal WD40 repeats and C-terminal coiled-coil domains (Fu et al., 2018; Urbanska et al., 2018), and FAP43 and FAP244 can functionally compensate for each other (Kubo et al., 2018). MOT7 (Kubo et al., 2018) shares some similarities with LAX28, but the sequence identity is too low to conclude homology with confidence. LAX28, TEX47 and MOT7 do not show clear signatures of known domains but may be able to adopt a similar 3D fold, as Phyre^2^ modelling predicted a beta-sheet with 4-5 strands and 2-3 alpha helices for all three proteins, revealing similarities to FAD-binding domains of the BLUF family. BLUF domains were first identified in *Euglena* (Iseki et al., 2002) and bacteria (Masuda and Bauer, 2002) and have been linked to sensing blue-light (Gomelsky and Klug, 2002). Interestingly, the blue-light sensing protein identified in *Euglena gracilis* has been characterised as a photoactivated adenylyl cyclase, containing two BLUF domains, that regulates flagellar motility by controlling intraflagellar levels of cyclic AMP (Iseki et al., 2002). A role in light-sensing for MOT7 seems worth exploring, since *Chlamydomonas* clearly undergoes phototaxis (Bennett and Golestanian, 2015). While *Leishmania* is not known to respond to light, BLUF domains may also function in redox-dependent signal transduction (Gomelsky and Klug, 2002). Thus, whether LAX28 is required only for IDA assembly or whether it contributes more directly to its regulatory role in the axoneme remains to be studied.

Given the role of IDA f/I1 as a regulatory hub for the modulation of flagellar waveforms one might perhaps predict that this is precisely where cell-type or species-specific proteins are required. Their role might be to orchestrate the conserved basic mechanisms of flagellar bend generation in biologically appropriate ways to produce the observed diversity of beating patterns. Further work in this area will no doubt benefit from comparative studies between cell types with a range of different behaviours.

While the precise phylogenetic relationship, if any, between LAX28, TEX47 and MOT7 remains to be established, TEX47 should be followed up for a possible link to male infertility. Male infertility affects about 7% of men and the genetic cause for this disease has been identified for only 25%. While more than 3000 genes have been associated with spermatogenesis, less than 0.01% of these have been linked to infertility (reviewed in Neto et al. (2016)). Compound-heterozygous mutations and frameshift mutations in CFAP44 and CFAP43 have been connected to male infertility (Tang et al., 2017). While to our knowledge mutations in TEX47 have not yet been linked to male infertility, the predominant expression of TEX47 in human testis (Fagerberg et al., 2014) its presence in the human sperm tail proteome (Amaral et al., 2013) and the results presented in this study for LAX28 suggest it may have a role in the regulation of sperm motility.

## Materials and Methods

### Cell culture

Promastigote-forms of *L. mex Cas9 T7* (Beneke et al., 2017) (derived from *L. mexicana* WHO strain MNYC/BZ/62/M379) were grown at 28°C in M199 medium (Life Technologies) supplemented with 2.2 g/L NaHCO_3_, 0.005% haemin, 40 mM 4-(2-Hydroxyethyl)piperazine-1-ethanesulfonic acid (HEPES) pH 7.4 and 10% FCS. 50 μg/ml Nourseothricin Sulphate and 32 μg/ml Hygromycin B were included before transfection with pPLOT or pT cassettes. For selection and maintenance of genetically modified *L. mex Cas9 T7* lines, the relevant selection drugs were added to supplemented M199 medium as described in (Beneke et al., 2017).

### Salt extracted axonemes

A widely utilized isolation protocol for *T. brucei* flagellar skeletons (Robinson and Gull, 1991) was adapted for use in *L. mexicana*. Cells were first non-ionic detergent extracted in 1% octyl glucoside to yield whole cell cytoskeletons and then salt extracted on ice using 1 M NaCl to depolymerize subpellicular microtubules. To avoid proteolytic degradation, all procedures were performed on ice or at 4°C during centrifugations. 1·10^9^ *L. mexicana ΔCFAP44* mutants and *L. mex Cas9 T7 parental* cells were collected at 800*g* for 15 min and resuspended in 2.5 ml phosphate buffered saline (PBS), containing a protease inhibitor cocktail [final concentration, 1x Halt Protease Inhibitor Cocktail (Thermo Fisher, containing 1 mM AEBSF•HCl, 0.8 µM Aprotinin, 50 µM Bestatin, 15 µM E64, 20 µM Leupeptin, 10 µM Pepstatin A) supplemented with 500 μM Phenylmethylsulfonyl fluoride (PMSF) and 5 mM EDTA]. 50 µl of cell suspension was isolated and supplemented with 5 µl 20% SDS solution (Fig. 1 (i)). The remaining cell suspension was centrifuged again and resuspended in 2.5 ml 10 mM PIPES [10 mM NaCl, 10 mM piperazine-N,N′-bis(2-ethanesulfonic acid, 1 mM CaCl_2_, 1 mM MgCl_2_, 0.32 M sucrose, adjusted to pH 7.2], containing protease inhibitor cocktail and octylglucoside (1% (w/v) final conc.). Cells were left on ice for 5 min and centrifuged for 10 min at 2000*g*. 50 µl of supernatant was isolated and supplemented with 5 µl 20% SDS solution (Fig. 1 (s/n 1)). 1.25 ml cold 10 mM PIPES buffer, containing protease inhibitor cocktail, octylglucoside (1% (w/v) final conc.) and 1 M NaCl was added to the pellet and vortexed for 60 sec. After incubation for 30 min on ice, the solution was centrifuged at 12,200*g* for 20 min. 50 µl of supernatant was isolated and supplemented with 5 µl 20% SDS solution (Fig. 1 (s/n 2)). The pellet was resuspended in 1 ml cold 10 mM PIPES buffer, containing a protease inhibitor cocktail and 0.32 M sucrose. The sample was loaded on top of a sucrose-bed containing one layer of 10 mM PIPES with 33% w/v sucrose [10 mM NaCl, 10 mM piperazine-N,N′-bis(2-ethanesulfonic acid, 1 mM CaCl_2_, 1 mM MgCl_2_, adjusted to pH 7.2 with 0.96M sucrose] and centrifuged at 800*g* for 15 min. While the top layer was centrifuged again at 12,200*g* for 20 min the remaining sucrose-bed was discarded. After centrifugation the supernatant was discarded and the pellet resuspended in 100 µl PBS containing protease inhibitor cocktail and 2% SDS (Fig. 1 (iv)). Proteins were quantified using BCA assay. For Coomassie gels samples were mixed with 4x Laemmli buffer (1x final concentration) and heated at 60°C for 10 minutes. For proteomic analysis ∼4 µg of protein (∼50 µl) of the final fraction (iv) were analysed by liquid tandem mass spectrometry.

### Proteomics

Protein samples were prepared using filter-aided sample preparation (FASP) digestion (Wisniewski et al., 2009) (Vicacon500, Sartorius, VN01H02 10kDA). The FASP filter were washed with 200 µl 0.1 % trifluoroacetic acid (TFA) in 50 % acetonitrile (ACN) (14,300*g*, 10min) and ∼4 µg of protein was loaded onto the filter. Samples were denatured with 200 µl 8 M urea in 100 mM TEAB (triethylammonium bicarbonate) for 30 min at RT, reduced with 10 mM TCEP (Tris(2-carboxyethyl)phosphine hydrochloride) for 30 min at RT and alkylated with 50 mM CAA (chloroacetamide) for 30min at RT in the dark. FASP columns were centrifuged (14,300*g*, 10min) and washed first multiple times with 200 µl 6 M urea in 50 mM TEAB until no more bubbles form and then washed twice with 200 µl 1 M urea in 50 mM TEAB. FASP columns were centrifuged again. Samples were digested with 200 ng trypsin in 300 µl 50 mM TEAB overnight at 37°C. Columns were centrifuged and flow through was kept. Columns were washed with 200 µl 0.1 % TFA and with 200 µl 50 % ACN in 0.1 % TFA. The flow through was collected from both washes. Samples were dried in a SpeedVac, resuspend in 50 µl 5% formic acid and 5% DMSO and then trapped on a C18 PepMap100 pre-column (300µm i.d. x 5mm, 100 Å, Thermo Fisher Scientific) using solvent A (0.1% Formic Acid in water) at a pressure of 500 bar and separated on an Ultimate 3000 UHPLC system (Thermo Fischer Scientific) coupled to a QExactive mass spectrometer (Thermo Fischer Scientific). The peptides were separated on an in-house packed analytical column (360µm x 75µm i.d. packed with ReproSil-Pur 120 C18-AQ, 1.9µm,120 Å, Dr. Maisch GmbH) and then electro sprayed directly into an QExactive mass spectrometer (Thermo Fischer Scientific) through an EASY-Spray nano-electrospray ion source (Thermo Fischer Scientific) using a linear gradient (length: 60 minutes, 15% to 38% solvent B (0.1% formic acid in acetonitrile), flow rate: 200 nL/min). The raw data was acquired on the mass spectrometer in a data-dependent mode (DDA). Full scan MS spectra were acquired in the Orbitrap (scan range 350-2000 m/z, resolution 70000, AGC target 3e6, maximum injection time 50 ms). After the MS scans, the 20 most intense peaks were selected for HCD fragmentation at 30% of normalised collision energy. HCD spectra were also acquired in the Orbitrap (resolution 17500, AGC target 5e4, maximum injection time 120 ms) with first fixed mass at 180 m/z.

### Proteomic analysis

MS-data were analysed as previously described (Beneke et al., 2019). Briefly, MS-data was converted from .RAW to .MGF file using ProteoWizard and uploaded to the Central Proteomics Facilities Pipeline (CPFP (Trudgian et al.)). Protein lists were generated by using CPFP meta-searches against the predicted *L. mexicana* proteome (gene models based on (Fiebig et al., 2015)), followed by label-free SINQ quantification (S1 Table). Proteins that were exclusively identified in *ΔCFAP44* mutants were generally of low abundance (2 or 3 detected peptides; S1 and S2 Table), with the exception of beta tubulin (LmxM.08.1230) for which 48 detected peptides were recorded in*ΔCFAP44* mutants. However, only 1 out of those 48 peptides was unique (spectral count 1; S1 and S2 Table). This suggests that proteins exclusively identified in the *ΔCFAP44* mutants may represent false positives and they were not further analysed in this study. The mass spectrometry proteomics data have been deposited to the ProteomeXchange Consortium via the PRIDE (Vizcaino et al., 2016) partner repository with the dataset identifier PXD014077.

### Solubilisation experiments

Solubilisation of CFAP44 was tested using a reporter cell line, expressing *in situ* tagged *CFAP44::eYFP* fusion protein. 1·10^7^ cells were collected at 800*g* for 5 min and washed once in PBS. Cells were pelleted again and resuspended in 100 µl of 10 mM PIPES buffer (as above), containing a protease inhibitor cocktail (as above) and octylglucoside (1% (w/v) final conc.), as well as either 2 M LiCl, 2 M CaCl_2_, 3.2 M KCl or 4 M NaCl. Lower concentrations were also tested including: 0.01 - 0.75 M NaCl, 1 M LiCl or 0.25 - 1.5 M CaCl_2_. Cell suspensions were vortexed for 60 sec and incubated for 30 min on ice. The solution was centrifuged at 17,000*g* for 2 min and the pellet washed once in PBS. Cells were pelleted again, resuspended in 10 mM PIPES buffer and pipetted onto a glass slide for viewing under a microscope.

### CRISPR-Cas9 gene knockouts and tags

Gene deletion and tagging was essentially done as described in Beneke et al. (2017). The online primer design tool www.LeishGEdit.net was used to design primers for amplification of the 5’ and 3’ sgRNA templates and for amplification of donor DNA from pT and pPLOT plasmids. Following transfection with pPLOT-cassettes, limiting dilution was used to generate clonal tagged cell lines (Beneke and Gluenz, 2019). These were then subjected to gene deletions using two different pT cassettes and selected as populations.

### Addback construction

The ORF of LAX28 was amplified using primers F: 5’-TTAGCAACTAGTATGGAACAGAAGCTGATCAGCGAAGAAGACCTGGAGCAAAAGCTCATTAGCGAGGAGGACCTCATGCCTCCGCGCAAGTCAGA-3’ and R: 5’-TTAGCACCATGGGCGCGGGTTCATTTTATCGT-3’ and cloned into pTadd (Beneke et al., 2017) using *Spe* I and *Nco* I cloning sites. 5 µg of circular plasmid was transfected as described previously (Beneke et al., 2017) to allow episomal expression of *2xMyc::LAX28*. Drug resistant cells were selected using 25 µg/ml phleomycin.

### Diagnostic PCR for knockout verification

Extracted genomic DNA of drug-selected populations was subjected to diagnostic PCRs to test for the presence of the target gene ORF in putative KO lines and the parental cell line as described in (Beneke and Gluenz, 2019; Beneke et al., 2017) and using primer sequences reported in Beneke et al. (2019). To show presence of genomic DNA in the test samples, a second PCR reaction was performed using primers F: 5’-CGCAGAAGGAGAAGAGCGAG-3’ and R: 5’-GTTGTACACGGACAGCTCCA-3’ to amplify the ORF of PFR2.

### Light and electron microscopy

*L. mexicana* expressing fluorescent fusion proteins were prepared as described in (Wheeler et al., 2015) and immediately imaged live, with a Zeiss Axioimager.Z2 microscope with a 63× numerical aperture (NA) 1.40 oil immersion objective and a Hamamatsu ORCA-Flash4.0 camera at the ambient temperature of 25–28°C. Micrographs were taken with a 3,000 ms exposure time for the green fluorescent channel.

For transmission electron microscopy, cells were prepared with a chemical fixation protocol similar to (Hoog et al.). Briefly, cells were fixed with 2.5% glutaraldehyde and 4% paraformaldehyde in M199 culture medium for 2 hours at room temperature. Fixed cells were washed six times for 10 min in 0.1 M PIPES buffer at pH 7.2. Wash four of six was supplement with 50 mM glycine. Cells were embedded in 4% low melting point agarose and incubated in 1% osmium tetroxide and 1.5% potassium ferrocyanide in 0.1 M PIPES buffer at 4 °C for 1 hour in darkness. Samples were then washed five times with ddH_2_O for 5 min each time and stained with 0.5% uranyl acetate in darkness at 4°C overnight. Samples were dehydrated, embedded in epoxy resin, sectioned and on-section stained as described previously (Hoog et al., 2010). Electron micrographs were captured on a Tecnai 12 TEM (FEI) with an Ultrascan 1000 CCD camera (Gatan).

### Image processing

All micrographs were processed using Fiji (Schindelin et al., 2012). To allow comparison between the fluorescent signal in tagged cell lines, the same settings were used to display the green fluorescence channel. Channel settings were set for CFAP44::mNG tagged cell lines 7,000 to 20,000, for IC140::eYFP tagged cell lines 6,000 to 11,000 (except in Fig. 2 A, 17,000 to 22,000) and for LAX28::mNG tagged cell lines 5,000 to 12,000. Settings were identical for the tagged cell line and additional knockouts and addbacks on top of the tagged cell lines.

For average rotation of axonemes in TEM images, nine sections from each cell line were average rotated as previously described (Gadelha et al., 2006; Wheeler et al., 2015). The resulting images were aligned using the “Align image by line ROI” function in Fiji (Schindelin et al., 2012). A stack was generated from these aligned images and signals were averaged by using the “Z projection” function.

### Motility analysis

Motility analysis was performed as previously described in Beneke et al. (2019) using the method from Wheeler (2017). Directionality ([Velocity/Speed]) and mean speed was measured for each mutant from three samples taken from cell cultures at a density of approximately 6·10^6^ cells/ml. 5 μl of cell culture was placed on a glass slide in a 250-μm deep chamber covered with a # 1.5 cover slip and imaged using darkfield illumination with a 10× NA 0.3 objective and a Hamamatsu ORCA-Flash4.0 camera on a Zeiss Axioimager.Z2 microscope at the ambient temperature of 25–28°C.

### Illumina sequencing

*Leishmania* genomic DNA was prepared using the Illumina TruSeq Nano DNA Library kit according to the manufacturer’s instructions. The final sequencing pool was quantified by qPCR using the NEB Library Quant Kit and library size was determined using the Agilent High Sensitivity DNA Kit on a 2100 Bioanalyzer instrument. The final library was multiplexed with other sequencing samples and the Illumina sequencer was loaded with 1.8 pM. Sequencing was performed using a NextSeq 500/550 High Output Kit v2.5 (2×75 cycles, 6 and 8 cycles index read).

NextSeq raw files were de-multiplexed using bcl2fastq (Illumina) with 0 nt mismatch for indexes and assembled using the Burrows-Wheeler Aligner (Li and Durbin, 2009). Samtools (Li et al., 2009) was used for sorting and indexing bam files. Bam files were viewed using the IGV genome browser (Robinson et al., 2011).

## Acknowledgements

We like to thank Svenja Hester and Philip Charles (Central Proteomics Facility) for help with protein mass spectrometry, Errin Johnson and Raman Dhaliwal (Dunn School Bioimaging Facility) for assistance with electron microscopy and Keith Gull (University of Oxford) for sharing equipment funded by the Wellcome Trust.

