## Supplemental Figures for "LAX28 is required for assembly of the inner dynein arm l1 and tether/tether head complex in the *Leishmania* flagellum"

### Supporting Information

#### S1 Table. SINQ raw data

Table shows the raw output from the CPFP pipeline with label-free SINQ quantification. The data table contains all identified proteins (listed by GeneIDs) from detergent/salt extracted axonemes of *Δ CFAP44* mutants and *L. mex Cas9 T7 parental* cells. The relative enrichment across both fractions and proteins uniquely identified in each sample are listed. Listed is the spectral count (not normalized enrichment) and spectral index (normalized enrichment).

#### S2 Table. GeneID list of the detergent/salt extracted proteome (p > 0.95, number of peptides > 2)

GeneIDs identified in the SINQ analysis (S1 Table) were filtered (p > 0.95, number of peptides > 2) and listed with additional information.

#### S1 Figure. Solubilisation of CFAP44

**(A)** Overview of attempts to solubilize CFAP44 in *L. mexicana*. *CFAP44::eYFP* expressing cells were detergent and salt extracted using octylglucoside (1% (w/v) final conc.) and various salts, as indicated. **(B)** Micrographs show merged phase and Hoechst DNA stain (red) for each isolation stage (i and ii) depicted in (A). **(i)** *L. mexicana CFAP44::eYFP* cells before detergent and salt extraction and **(ii)** cells after detergent and salt extraction using either 2 M LiCl, 2 M $CaCl_2$, 3.2 M KCl or 4 M NaCl as indicated. Scale bars represent 10 µm.

#### S2 Figure. Null mutant verification by diagnostic PCR

**(A)** Strategy for amplification of a fragment of the target gene ORF, using ORF-internal primers (small arrows). **(B, C)** PCR amplicons run on an agarose gel, comparing results for genomic DNA from the putative knockout cell line (K) and the relevant parental cell line (P; *L. mex* Cas9 T7 *IC140::eYFP* in B or *L. mex* Cas9 T7 *LAX28::mNG* in C). Loss of the target OFR is indicated by absence of a band. Fainter bands below the target gene amplicon are likely primer dimers. The gene names below the image indicate the target gene ORF. See Beneke et al. (2019) S7 Table for amplicon sizes. Left-most lane: DNA ladder, size of bands in bp is indicated on the left. **(D)** Strategy for verifying presence of genomic DNA in the test samples, using *PFR2* ORF-internal primers (small arrows). **(E, F)** PCR amplicons for the *PFR2* ORF visualised on an agarose gel, and presence confirmed for all transfected cell lines as well as the parental cell line *L. mex* Cas9 T7 (Parental). Left-most lane: DNA ladder, size of bands in bp is indicated on the left.

**(G)** Strategy for validation of null mutants shown in Fig. 2 C, 2 D, 3 and S4: amplification of a fragment of the *LAX28* ORF, **(H)** amplification of a fragment of the *PFR2* ORF. **(I)** PCR amplicons of the *LAX28* ORF and of the *PFR2* ORF run on an agarose gel. Template DNA was extracted from the cell lines indicated below the image. Two independent populations of *ΔLAX28* mutants in the *CFAP44::mNG* background were produced, with indistinguishable phenotypes. Population 1 (POP1) was used for all experiments shown in this study.

### S3 Figure. Western blot analysis of fusion protein expression

Western blot detection of tagged proteins in different *L. mexicana* cell lines, as indicated. Whole cells were lysed in Laemmli buffer at 65°C for 10 min, and protein samples from $2 \cdot 10^6$ cells were separated on a 10% SDS PAGE gel, transferred to a nitrocellulose membrane and probed with anti-myc antibody 4A6 (Merck; 1:2000 dilution in TBST 5% (w/v) skimmed milk powder). For detection, an HRP-conjugated secondary antibody was used followed by detection of chemiluminescent substrate and exposure to X-ray film, suing standard procedures. All *mNG* and *eYFP* tags used in this study were fused to 3 copies of the myc epitope tag; the addback copy of *LAX28* was fused to 2 myc epitope tags. Arrows on the right indicate the expected fusion proteins detected by the myc antibody. Molecular weights (MW) in kDa are indicated on the left. Lower panel: image of the membrane stained with Ponceau red before antibody detection.

### S4 Figure. Detection of LAX28::mNG signal over the *Leishmania* cell cycle in *ΔIC140* and *ΔCFAP44* null mutants

Fluorescence micrographs showing *L. mexicana* cell lines expressing LAX28::mNG, *L. mexicana* LAX28::mNG carrying a *Δ CFAP44* deletion and *L. mexicana* LAX28::mNG carrying a *Δ IC140* deletion, Three different cell cycle stages are shown for each cell line, staged according to the number of kinetoplasts (K), nuclei (N) and Flagella (F). Left column: Merged phase and fluorescence channels; red, Hoechst-stained DNA, green, mNG or eYFP signal, respectively. Right column: grayscale rendition of green fluorescence channel. Scale bar, 5 µm.

### S5 Figure. Illumina sequencing of ***ΔLAX28*** null mutants

Whole genomic DNA from the parental cell line *L. mex* Cas9 T7 and the *ΔLAX28* deletion mutant was sequenced using Illumina NextSeq. **(A)** Sequence reads were viewed in the IGV genome browser (Robinson et al., 2011). The screenshot shows the LAX28 (LmxM.24.1310) locus on chromosome 24. Analysis of the read coverage for *ΔLAX28* shows there were almost zero reads spanning the *LAX28* ORF while the *LAX28* locus is well covered in *L. mex Cas9 T7* parental DNA. The single read pair that aligned to the *LAX28* locus in *ΔLAX28* is most likely to represent cross-contamination during genome sample preparation, as examination of the data excluded possible other technical explanations (e.g. demultiplexing error or poor read quality). **(B)** Blast queries using ORFs of *LAX28*, blasticidin , puromycin and neomycin resistance genes against sequence reads from *ΔLAX28* mutants and *L. mex Cas9 T7 parental* cells. Only LAX28 was detected in the parental DNA reads, while the blasticidin and puromycin resistance genes used to replace LAX28 were detected in the deletion mutants. Two reads of *LAX28* were identified in *ΔLAX28* mutants, corresponding to the read pair illustrated within the *LAX28* locus in (A).

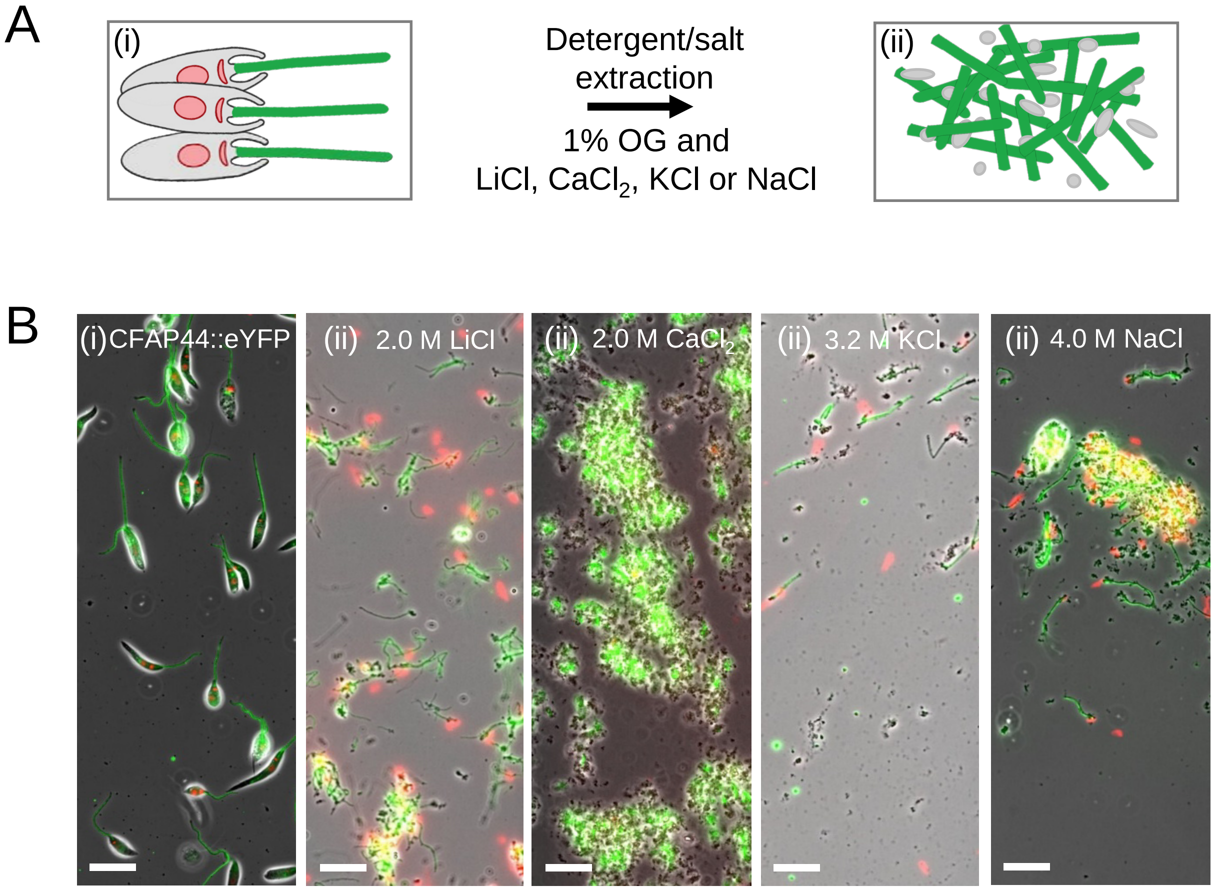
A
(i)
Detergent/salt
extraction
1% OG and
LiCl, CaCl2, KCl or NaCl
(ii)
B
(i) CFAP44::eYFP
(ii) 2.0 M LiCl
(ii) 2.0 M CaCl2
(ii) 3.2 M KCl
(ii) 4.0 M NaCl

S1 Figure

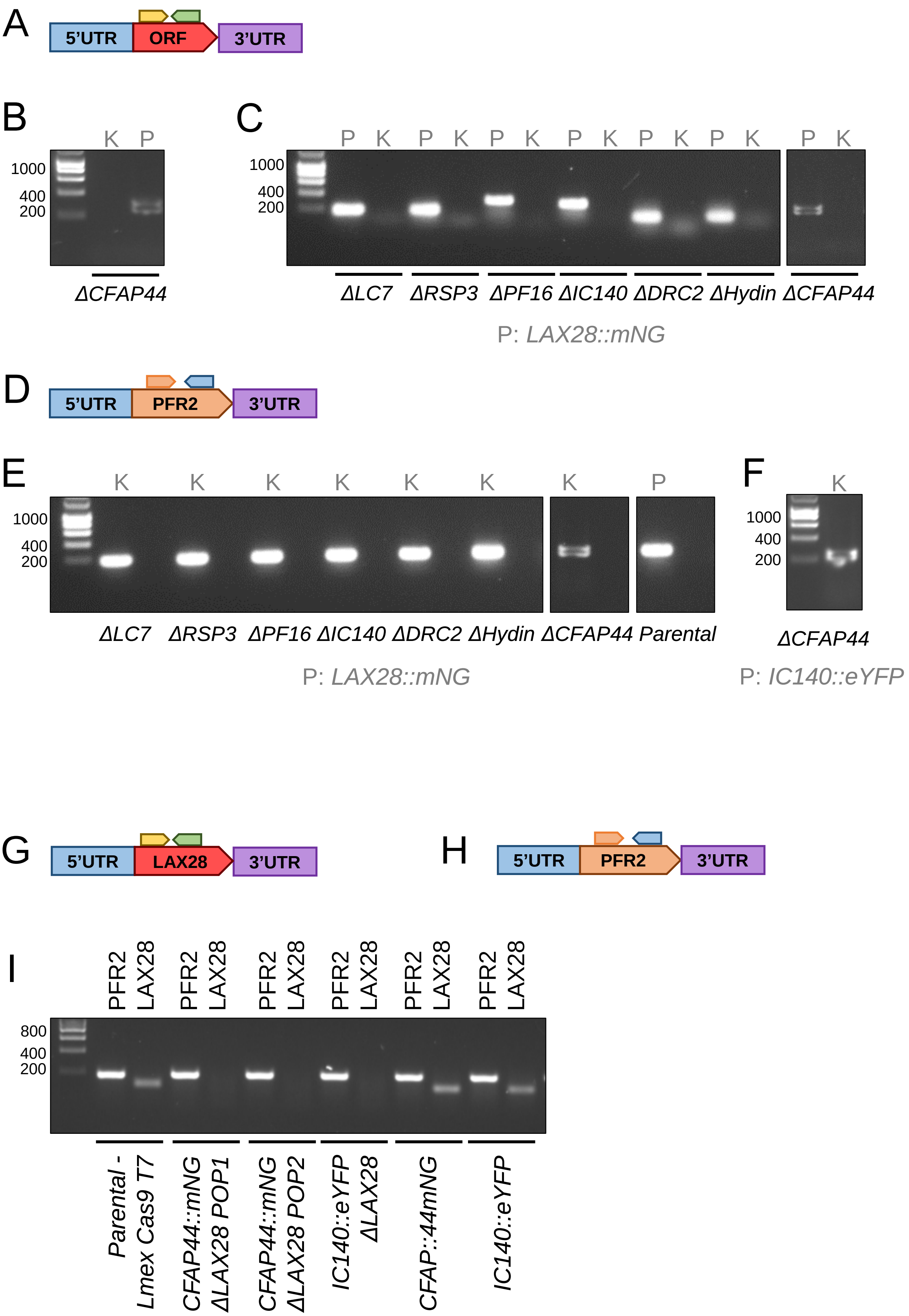
A
5'UTR
ORF
3'UTR
B
K P
1000
400
200
ΔCFAP44
C
P K P K P K P K P K P K P K
1000
400
200
ΔLC7 ΔRSP3 ΔPF16 ΔIC140 ΔDRC2 ΔHydin ΔCFAP44
P: LAX28::mNG
D
5'UTR
PFR2
3'UTR
E
K K K K K K K P
1000
400
200
ΔLC7 ΔRSP3 ΔPF16 ΔIC140 ΔDRC2 ΔHydin ΔCFAP44 Parental
P: LAX28::mNG
F
K
1000
400
200
ΔCFAP44
P: IC140::eYFP
G
5'UTR
LAX28
3'UTR
H
5'UTR
PFR2
3'UTR
I
PFR2 LAX28 PFR2 LAX28 PFR2 LAX28 PFR2 LAX28 PFR2 LAX28 PFR2 LAX28
800
400
200
Parental - Lmex Cas9 T7
CFAP44::mNG ΔLAX28 POP1
CFAP44::mNG ΔLAX28 POP2
IC140::eYFP ΔLAX28
CFAP::44mNG
IC140::eYFP

S2 Figure

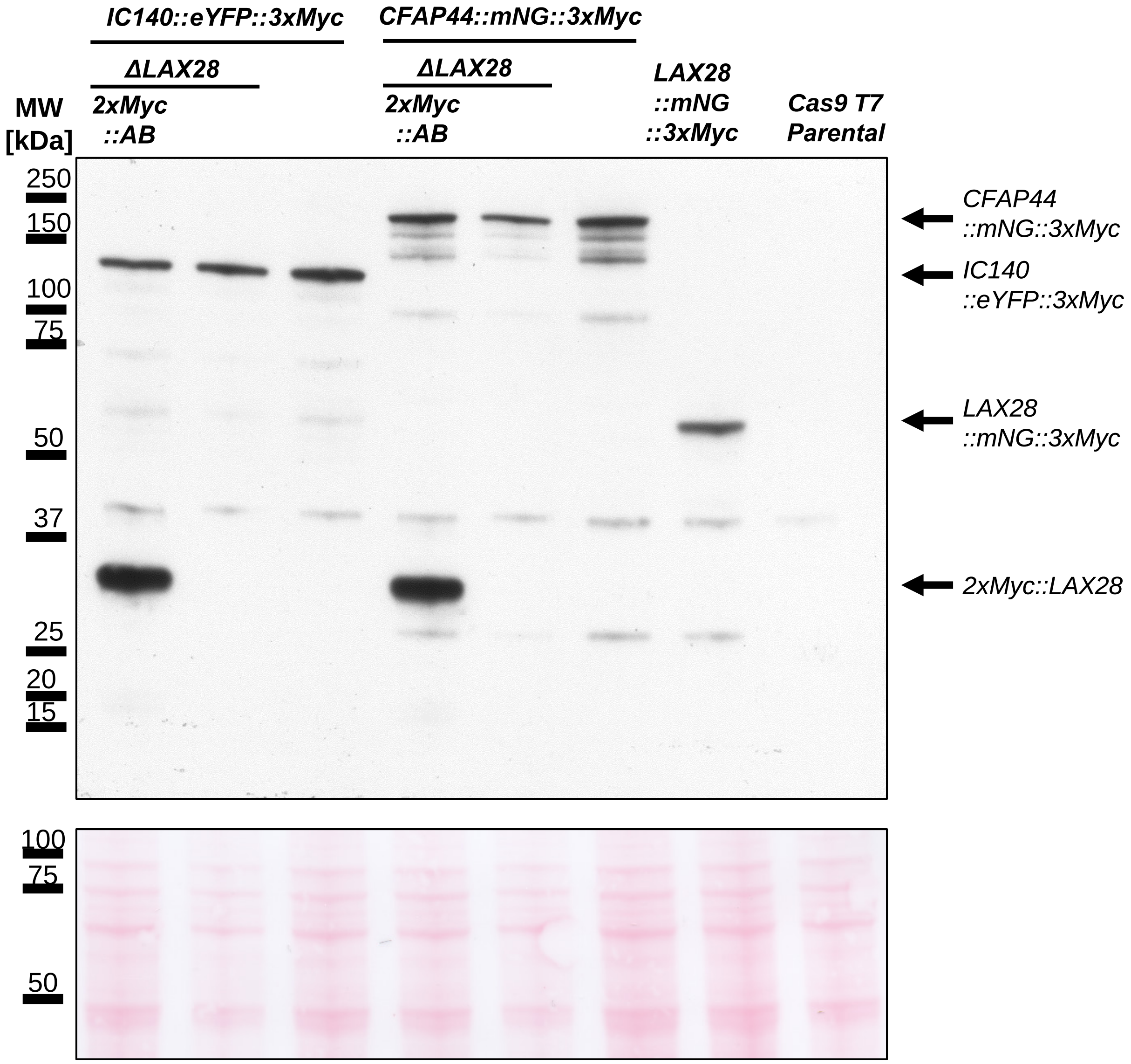

IC140::eYFP::3xMyc
CFAP44::mNG::3xMyc
ΔLAX28
ΔLAX28
MW
[kDa]
2xMyc
::AB
2xMyc
::AB
LAX28
::mNG
::3xMyc
Cas9 T7
Parental
250
150
100
75
50
37
25
20
15
CFAP44
::mNG::3xMyc
IC140
::eYFP::3xMyc
LAX28
::mNG::3xMyc
2xMyc::LAX28
100
75
50


S3 Figure

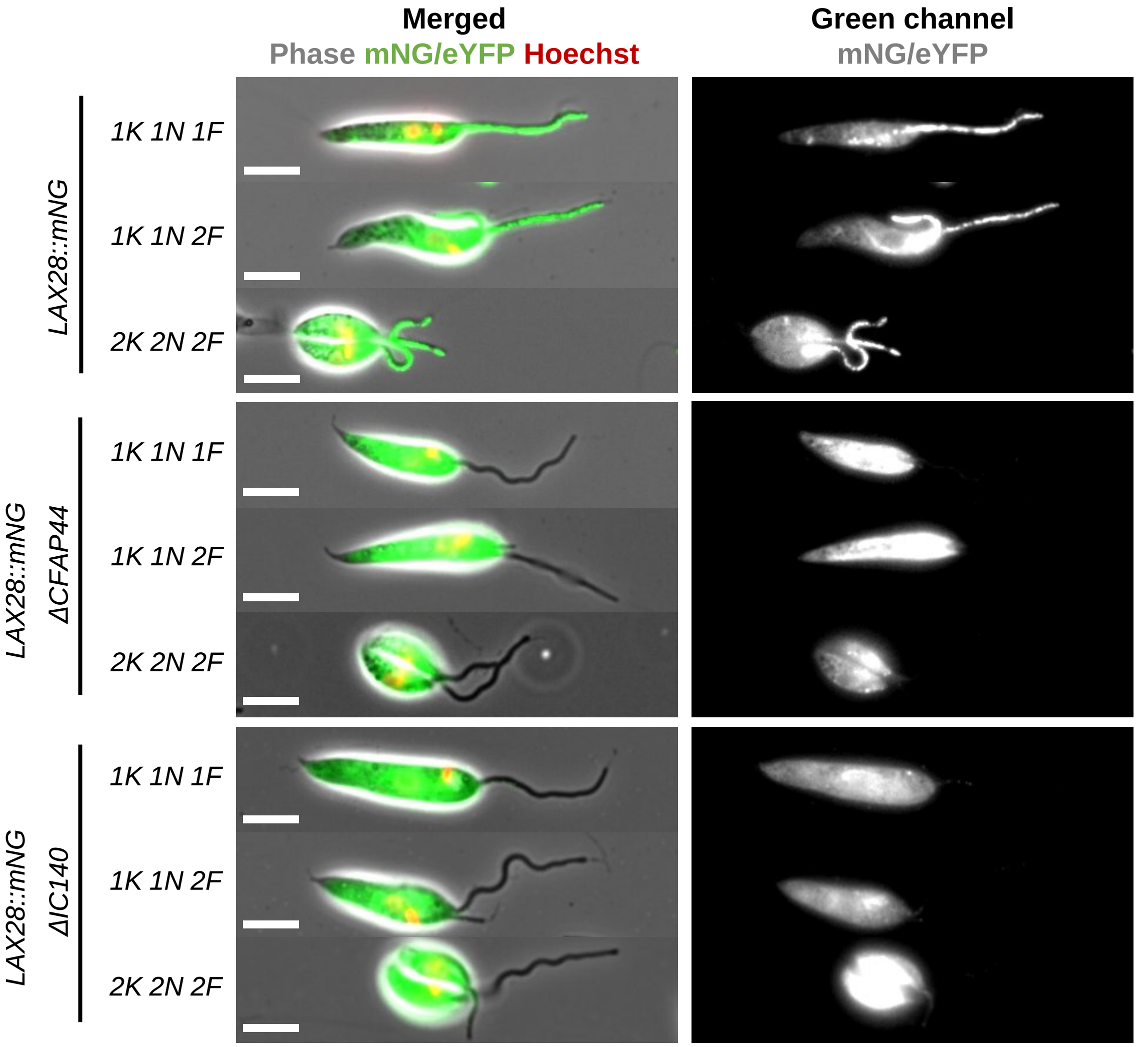

Merged
Phase mNG/eYFP Hoechst
Green channel
mNG/eYFP
LAX28::mNG
1K 1N 1F
1K 1N 2F
2K 2N 2F
LAX28::mNG ΔCFAP44
1K 1N 1F
1K 1N 2F
2K 2N 2F
LAX28::mNG ΔIC140
1K 1N 1F
1K 1N 2F
2K 2N 2F


S4 Figure

A

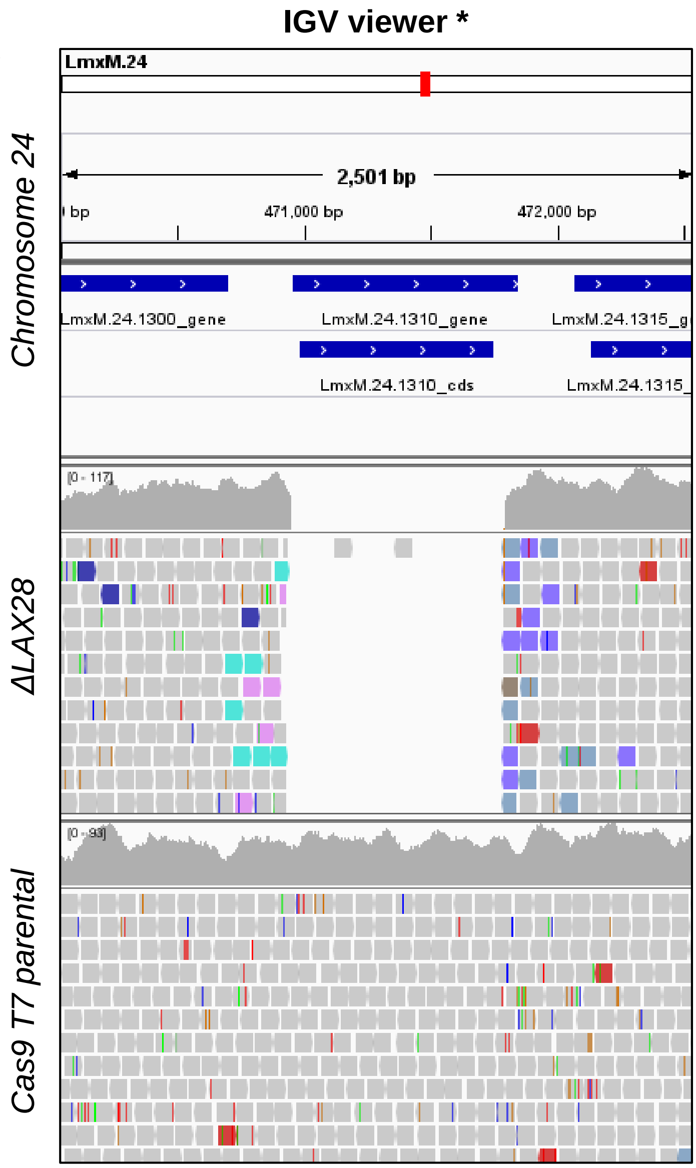


* Robinson et al., Nature 2011

B

| ORF blast query | Aligned number of reads to ORF query: *ΔLAX28* | Aligned number of reads to ORF query: *Cas9 T7 parental* |
|---|---|---|
| LAX28 | 2 | >750 |
| Blastacidin | 248 | 0 |
| Puromycin | 356 | 0 |
| Neomycin | 0 | 0 |

S5 Figure
